# HSP40 overexpression in pacemaker neurons protects against circadian dysfunction in a *Drosophila* model of Huntington’s Disease

**DOI:** 10.1101/2021.12.27.474320

**Authors:** Pavitra Prakash, Arpit Kumar Pradhan, Vasu Sheeba

**Affiliations:** Evolutionary and Integrative Biology Unit, Jawaharlal Nehru Centre for Advanced Scientific Research, Bangalore, India - 560064.; Neuroscience Unit, Jawaharlal Nehru Centre for Advanced Scientific Research, Bangalore, India - 560064.

**Keywords:** Circadian, Heat Shock Protein (HSP), HSP40, Huntingtin, Huntington’s Disease, Neurodegeneration, *Drosophila*, LNv

## Abstract

Circadian disturbances are early features of neurodegenerative diseases, including Huntington’s Disease (HD), affecting the quality of life of patients and caregivers. Emerging evidence suggests that circadian decline feeds-forward to neurodegenerative symptoms, exacerbating them, highlighting a need for restoring circadian health. Therefore, we asked whether any of the known neurotoxic modifiers can suppress circadian dysfunction. We performed a screen of neurotoxicity-modifier genes to suppress circadian behavioural arrhythmicity in a *Drosophila* circadian HD model. Notably, the molecular chaperones HSP40 and HSP70 (<u>H</u>eat <u>S</u>hock <u>P</u>rotein) emerged as significant suppressors in the circadian context, with HSP40 being the more potent mitigator of HD-induced deficits. Upon HSP40 overexpression in the *Drosophila* circadian ventrolateral neurons (LNv), the behavioural rhythm rescue was associated with neuronal rescue of loss in circadian proteins from small LNv soma. Specifically, there was a restoration of the molecular clock protein Period and its oscillations in young flies and a long-lasting rescue of the output neuropeptide Pigment Dispersing Factor. Significantly, there was a reduction in the expanded Huntingtin inclusion load, concomitant with the appearance of a spot-like Huntingtin form. Thus, we provide evidence for the first time that implicates the neuroprotective chaperone HSP40 in *circadian rehabilitation*. Given the importance of proteostasis and circadian health in neurodegenerative diseases, the involvement of molecular chaperones in circadian maintenance has broader therapeutic implications.

**Summary Statement:** This study shows, for the first time, a neuroprotective role of chaperone HSP40 in overcoming circadian dysfunction associated with Huntington’s Disease in a *Drosophila* model

## Introduction

Huntington’s disease (HD) is a neurodegenerative disease (ND) due to a dominant mutation in the huntingtin gene (*Htt*), leading to the expansion of the glutamine (Q) amino acid repeat tract in the Huntingtin protein (HTT) beyond a threshold of 35-40 poly Glutamine (polyQ) repeats. It shares several features with other NDs (Gusella and MacDonald, 2006; Bates et al., 2015), such as a typical middle-age onset, specificity of brain areas affected, motor and cognitive impairments, psychiatric disturbances, a progressive worsening with age, decline in the quality of life and longevity. The presence of disease protein aggregates, aberrant proteostasis, synaptic toxicity, oxidative stress and neurodegeneration are shared pathophysiological mechanisms.

Circadian and sleep disruptions are now recognised as early symptoms of many NDs, including HD (Goodman et al., 2011; Morton et al., 2014; Lebreton et al., 2015; Bellosta Diago et al., 2017). The mammalian clock centre is affected in HD mice, including molecular clock disruptions and reduction in the Vasoactive Intestinal Peptide, a clock output neuropeptide (Morton et al., 2005; Maywood et al., 2010; Kudo et al., 2011; van Wamelen et al., 2013). Emerging evidence supports bi-directional crosstalk between the circadian and neurodegenerative axes, with circadian function impacting the aetiology and progression of NDs (Hood and Amir, 2017; Leng et al., 2019; Carter et al., 2021; Voysey et al., 2021). Improving clock function and sleep in HD mice have been neuroprotective (Pallier et al., 2007; Maywood et al., 2010; Ouk et al., 2017; Wang et al., 2017; Whittaker et al., 2018), whereas clock disruptions worsen ND (Krishnan et al., 2012; Lauretti et al., 2017; Kim et al., 2018; Sharma and Goyal, 2020).

Given the beneficial effects of circadian improvement on neurodegeneration, we aimed to uncover modifiers of expHTT-induced circadian arrhythmicity that could also serve as modifiers of neurodegenerative phenotypes. We have previously established and characterised a circadian model of HD in *Drosophila melanogaster* (Sheeba et al., 2008; Sheeba et al., 2010; Prakash et al., 2017); flies expressing an expanded HTT (expHTT) with 128 polyQ (HTT-Q128) in a subset of the pacemaker neurons, the ventral lateral neurons (LNv). The LNv express a critical circadian output neuropeptide, the Pigment Dispersing Factor (PDF) and are comprised of the ∼4 small LNv (sLNv) and 4-5 large LNv (lLNv) (Helfrich-Förster, 1995; Renn et al., 1999). PDF and the sLNv are essential for locomotor activity/rest rhythms in constant darkness (DD) (Renn et al., 1999; Grima et al., 2004; Stoleru et al., 2004; Shafer and Taghert, 2009). Most flies expressing expHTT in the PDF^+^ LNv (*pdf>Q128*) had disrupted behavioural activity rhythms in DD, a loss of PDF from sLNv soma, loss of PER and its oscillations from LNv and presence of expHTT inclusions (Sheeba et al., 2008; Prakash et al., 2017). In these *pdf>Q128* flies, we carried out a genetic screen for modifiers that can rescue circadian behavioural arrhythmicity. These genes are grouped under different categories based on function (Table S1), and the expressed proteins assist in neuronal function and are modifiers of neurodegeneration (Steffan et al., 2001; Gunawardena et al., 2003; Shulman and Feany, 2003; Sang et al., 2005; Wyttenbach and Arrigo, 2009; Zhang et al., 2009; Sutton et al., 2013; Kampinga and Bergink, 2016; Menzies et al., 2017; Metaxakis et al., 2018). Co-expression of expHTT in LNv with candidates from the Heat Shock Protein (HSP) or autophagy group significantly improved the rhythmicity of flies than their expHTT-only expressing counterparts (Table S1). In the HSP group, these included *HSP23*, *HSP40* and *HSP70* homologs. The *tim>Q128,HSP70* also had better rhythmicity than *tim>Q128*. The central heat shock protein HSP70 and the co-chaperone HSP40 were chosen for further analysis.

HSPs play a central role in cellular proteostasis, aiding in protein folding, trafficking, degradation, preventing aberrant interactions, and disaggregation (Hartl and Hayer-Hartl, 2009; Kim et al., 2013; Nillegoda et al., 2018). HSP40 and HSP70 co-localise with mutant HTT aggregates (Jana, 2000; Muchowski et al., 2000; Kim et al., 2002; Scior et al., 2018) and their levels reduce with age, coinciding with proteostasis decline and middle-age HD onset (Hay, 2004; BenZvi et al., 2009; Taylor and Dillin, 2011; Brehme et al., 2014; Hipp et al., 2019). HSP upregulation alleviates proteotoxic stress and HD symptoms (Jana, 2000; Hansson et al., 2003; Branco et al., 2008; Labbadia et al., 2012; Brehme and Voisine, 2016; Kakkar et al., 2016), while their reduction aggravates neurodegenerative phenotypes in cell culture and animal models (Tagawa et al.,; Hageman et al., 2010; Jiang et al., 2012; Scior et al., 2018 Therefore, the emergence of HSPs as modifiers in our screen is not surprising. However, the role of HSPs in circadian rehabilitation is relatively unexplored.

We investigated the role of HSP40 and HSP70 as modifiers of expHTT-induced circadian neurodegenerative phenotypes in *Drosophila*. Of the two HSPs, HSP40 emerged as the more potent modifier of expHTT-induced phenotypes. Its overexpression rescued rhythmic locomotion over a substantial duration, PDF loss from sLNv, restored PER cycling in sLNv of young flies and decreased visible expHTT inclusion load favouring a new feature - a spot-like form of expHTT. HSP70 overexpression rescued rhythmicity and lowered expHTT inclusion numbers only at an early age, without rescuing PDF or PER in sLNv or affecting inclusions as the predominant form of expHTT in LNv. Co-expression of HSP40 and HSP70 in *pdf>Q128* leads to a synergistic improvement in the consolidation of activity rhythms. Overall, the present study establishes a role for HSPs as suppressors of expHTT-induced circadian impairments in a *Drosophila* circadian HD model.

## Materials and Methods

### Fly lines

The *UAS-HTTpolyQ* lines used in this study, namely *w;+;UAS-HTT-Q0A;+* (without Q repeats) and *w;+;UAS-HTT-Q128C;+* (with 128 polyQ repeats), were a gift from Troy Lee et al., 200). They contain the first 548 amino acids of the human *Htt* gene. The *UAS* males were crossed with virgin females of either *w;pdfGal4:+* or *w^1118^:+:+* (BL 5905) to generate fly lines expressing HTT-polyQ in the PDF^+^ LNV neurons designated *pdf>Q0* and *pdf>Q128* or the *UAS* background controls designated *Q0* and *Q128*, respectively. In some instances, a broader circadian driver, the *w;timGal4:UAS-GFP* was used for the screen. For the genetic screen, the fly lines used are listed in Table S1. The genes were either overexpressed or downregulated in the *pdf>Q128* background (and in a few cases *tim>Q128* background) (Table S1). A recombinant line of *w;pdfGal4;+* and *w;UAS-HTTQ128/+;+* was generated, denoted by *w;pdf-Q128;+* and used for the screen. The sample size for the screen varied between 16 to 32 flies per genotype.

For most of this study, the fly lines of focus were generated using the two HSP *UAS* lines, namely *w;+;UAS-DnaJ1k* (Bl 30553) and *w;+;UAS-HSPA1L* (Bl 7054). The human HSP70 homolog used in this study was HSPA1L (Bl 7454) (https://flybase.org/reports/FBgn0029163.html), as a recent analysis revealed that among the various chaperone families, the HSP70 family most frequently provided considerable proteotoxic relief in a variety of protein misfolding diseases (Brehme and Voisine, 2016). The next potent proteotoxic suppressor was the HSP40 family, among which DNAJB4 and DNAJB6 were the most potent polyQ disease modifiers (Brehme and Voisine, 2016). The dHDJ1 (DnaJ1k or CG10578 or DnaJ-1 or Dm HSP40) (Bl 30553) used in this study is a *Drosophila* ortholog of members of the human HSP40 Class B family with varying degrees of homology (DNAJB5, DNAJB4 and DNAJB1) (https://flybase.org/reports/FBgn0263106.html).

The *UAS-HSP* lines were used to generate the *w;pdfGal4/UAS-Q128;UAS-DnaJ1k/+* or *w;pdfGal4/UAS-Q128;HSPA1L/+* lines which would respectively overexpress HSP40 or HSP70 in the PDF^+^ LNv neurons, also expressing HTT-Q128. The above-generated lines are referred to as *pdf>Q128,HSP40* and *pdf>Q128,HSP70* throughout the text. Their *UAS* control lines are referred to as *Q128,HSP40* and *Q128,HSP70* and their *driver* control lines as *pdf>HSP40* and *pdf>HSP70*. Their corresponding *Q0* lines are *pdf>Q0,HSP40* and *pdf>Q0,HSP70*. All other relevant background controls were also used. Due to space constraints, in some figures (Figs 6A, C, S3B, C), *pdf>Q128*, *pdf>Q128,HSP40* and *pdf>Q128,HSP70* are abbreviated as *Q128*, *H40* and *H70*, respectively. Flies co-expressing both HSP40 and HSP70 along with HTT-Q128 in the LNv are referred to as *pdf>Q128,HSP40,HSP70* and their *Q0* counterparts are *pdf>Q0,HSP40,HSP70*. Crosses were maintained under 12 h:12 h light: dark cycles (LD), with ∼200 lux intensity of light phase, at 25. Flies were moved to DD 25 after two days post-eclosion for behavioural and immunocytochemical assays. All flies and crosses were maintained on a standard cornmeal medium.

### Behavioural assays

Most of the locomotor activity setup, assay conditions and analyses performed are described in a previous study (Prakash et al., 2017). Briefly, the activity rhythms of 3d virgin male flies were recorded in 7 mm glass tubes using the *Drosophila* Activity Monitoring system 2 (DAM2) from TriKinetics (TriKinetics, Waltham, MA). Activity recordings were carried out in an incubator (Percival DR36VL) at 25°C in constant darkness (DD) for at least 21 days (age 3d to 23d) with about 80% humidity. The data thus obtained were analysed using the Chi-Square periodogram in the CLOCKLAB software (Actimetrics, Wilmette, IL). The periodogram threshold was set at *p* = 0.01 (Pfeiffenberger et al., 2010). A fly was considered rhythmic if its periodogram amplitude was above the threshold and confirmed by visual inspection of the actogram by a single-blind analyser. The various activity rhythm features such as rhythmicity, rhythm strength or rhythm robustness and period were calculated over three 7d windows to view progressive changes with age. The three temporal windows (age windows or AWs) spanning 21d thus obtained are designated age window 1 (AW1: age 3d- 9d), age window 2 (AW2: age 10 d-16d) and age window 3 (AW3: age 17d -23d).

Additionally, to track daily changes in activity/rest rhythms, the extent of activity consolidation ‘*r*’ was also calculated using a MATLAB code as previously described (Prakash et al., 2017) with a few modifications. ‘*r*’ represents the extent to which activity data-points are consolidated over a circadian cycle of activity. The activity time series for a genotype was obtained at a resolution of 20 min bins. This time series was divided into cycles of length determined by the period (1) of that genotype, thereby identifying each circadian cycle in the time series. Thus, each ‘day’ used to calculate ‘daily’ r was obtained as a modulo T value. On each ‘day’, activity counts were imagined as unit vectors whose direction represented the time point (t) at which the count occurred within the cycle. Since counts were clustered into 20 min intervals, the data can be represented as vectors with a constant angular separation of 20*2π/T rad and magnitudes corresponding to the number of activity counts in each interval. *R* was calculated as the magnitude of the mean of these activity vectors. The rectangular coordinates of the mean vector were obtained as follows (Zar, 2010) – X = ΣA_t._cosθ _t_ /Σ A_t_ and Y= Σ A_t_sin _t_ θ_t_/ΣA_t_, where A_t_ represents activity counts at a given time point t and _t_ represents the vector angle associated with the timepoint. The magnitude of this vector ‘*r*’ was calculated as √X^2^ + Y^2^. Greater the magnitude of *r*, better is the degree of consolidation, with most of the activity occurring over a few closely spaced time-points. Lower the magnitude of *r*, the activity is spread over time and poorly consolidated. Given that, sometimes period changes were observed across age for a fly, daily cycles were identified separately for each 7d AW using the mean period values for the corresponding AW. For arrhythmic flies, the cycle length was determined using the mean period of the genotype for that AW. For a fly dying in the middle of an AW, the period of that fly in the prior AW (if any) was used to determine cycle length. The daily ‘*r*’ was averaged across flies to obtain the mean daily ‘*r*’.

At least three independent activity runs were carried out for overexpression of HSP40 with its *Q0* and *UAS* controls, all giving similar results. For overexpression of HSP70, three independent activity runs were carried out with its *Q0* controls, giving similar results. At least one experiment was carried out with all possible genetic controls for both sets of HSP40 and HSP70 overexpression experiments. No statistical tests were carried out to determine the minimum required sample sizes. However, as recommended (ostadinov et al., 2021), one full DAM2 monitor accommodating 32 flies per genotype was set up at the start of the experiment. The average percentage rhythmicity across multiple runs is plotted for HSP40 and HSP70 overexpression experiments. In addition, the percentage rhythmicity, period, robustness, and extent of activity consolidation ‘*r*’ values of a representative run with all relevant controls for each of the above overexpression experiments are plotted. For synergistic effects, a single run was carried out. There were fly deaths in AW3; therefore, AW3 analyses had fewer samples. When the fly numbers went below 10 (e.g. a few instances in AW3), those genotypes were excluded from statistical analyses for that AW. In cases where very few flies (n<10) were rhythmic like *pdf>Q128* number across AWs in the HSP70 overexpression and synergistic effect experiments and during AW3 in HSP40 overexpression experiment or *pdf>Q128,HSP40* during AW3 in the HSP40 overexpression experiment or most of the experimental genotypes during AW3 in the synergistic effect experiment, those genotypes were excluded from the between-genotype statistical analyses of period and rhythm robustness for that AW. Indicated in Table S2 are the numbers of surviving flies in AW2 and AW3 across experiments.

### Statistical analysis of activity rhythms

Data sets were first tested for normality using Shapiro-Wilk’s test and then for variance homogeneity using Levene’s test. Across experiments, the data comparing period or activity consolidation ‘*r*’ between genotypes for an AW or age did not satisfy the ANOVA assumption of normality, despite transformations. So was the case for rhythm robustness in the HSP40 overexpression experiment. Therefore, the non-parametric Kruskal-Wallis test of ranks followed by multiple comparisons of mean ranks was used. For the HSP70 overexpression experiment, datasets comparing rhythm robustness between genotypes for an AW were normally distributed post-transformation, but variances were not always homogenous. Hence, a Welch’s ANOVA was used, followed by Games-Howell post-hoc test. For the synergistic effect experiment, the data comparing robustness between genotypes for an AW was normally distributed and their variances homogenous. A one-way ANOVA followed by unequal N Tukey’s HSD tests were carried out. Friedman’s test for repeated measures was used to compare robustness, period and ‘*r*’ between AWs or ages for a genotype. Then Wilcoxon matched-pairs tests (or Conover Test for ‘*r*’) with Bonferroni correction (or Benjamini-Hochberg (BH) procedure to decrease the False Discovery Rate (FDR) for ‘*r*’; FDR set at 5%) on the pair-wise *p* values were used. A m x n Fisher’s Exact test, followed by multiple 2×2 Fisher’s Exact tests with BH procedure on all relevant comparisons, were used (using R) to compare the proportions of rhythmic flies between genotypes for an AW. Cochran Q test on the dichotomous variable rhythmicity (rhythmic and arrhythmic categories) was used to compare the proportion of rhythmic flies between AWs for a genotype, followed by multiple 2×2 McNemar’s tests on the dependent samples and BH procedure on all relevant comparisons. For comparing the mean rhythmicity of multiple independent runs between genotypes or between age, a repeated-measures ANOVA followed by Tukey’s HSD was performed post-arcsine conversion of the square-root transformed data.

Statistical analyses were executed using STATISTICA^TM^ 7.0 (StatSoftInc, 2004) and R (RCoreTeam, 2013). Welch’s ANOVA was performed using a Microsoft Excel template from http://www.biostathandbook.com/onewayanova.html (McDonald, 2014), McNemar’s test using SciStatCalc (https://scistatcalc.blogspot.com/2013/11/mcnemars-test-calculator.html) and Friedman’s test followed by Conover test for ‘*r*’ using ASTATSA (https://astatsa.com/FriedmanTest/).

### Immunocytochemical assays

The dissections and immunocytochemistry procedures performed were as described previously (Prakash et al., 2017). Adult fly brain tissues were dissected in 1X PBS at different ages, fixed with 4% paraformaldehyde at room temperature, blocked and stained with the appropriate primary antibodies, followed by secondary antibodies, and then mounted on slides using 70% glycerol in 1XPBS. Primary antibodies used for double staining were anti-HTT mouse (1:500) (MilliporeSigma MAB2166) and anti-PDF rabbit (1:30,000) (Michael Nitabach, Yale University) (Nitabach et al., 2006) and for triple staining, anti-PER rabbit (1:20,000) (Jeffrey C Hall, Brandeis University) (Houl et al., 2008), anti-HTT mouse (1:500) and anti-PDF rat (1:3000) (Jae Park, Vanderbilt University) (Park et al., 2000). Rabbit anti-PER was pre-absorbed onto *per^01^* embryos at 1:100 dilution. Secondary antibodies (1:3000) used were from Invitrogen: Alexa Fluor anti-rabbit 488, anti-mouse 546, anti-mouse 647 and anti-rat 555.

For characterising the PER^+^ and PDF^+^ LNv soma numbers, adult fly brain dissections were carried out in parallel at different ages. Since, with *pdf>Q128,HSP40*, a sustained rescue for two AWs was seen in behaviour, dissections were carried out in 3d, 9d, and 16d old flies corresponding to the beginning of AW1, the transition of AW1-AW2 and end of AW2, respectively. With *pdf>Q128,HSP70*, rhythm rescue was restricted to AW1. Hence, dissections were carried out at 3d and 9d, corresponding to the beginning and end of AW1, respectively. For characterising HTT status in LNv, adult fly brain dissections were carried out in parallel for the five genotypes (*pdf>Q128*, *pdf>Q0,HSP40*, *pdf>Q128,HSP40*, *pdf>Q0,HSP70* and *pdf>Q128,HSP70*) at 3d and 9d. Many of the HSP expressing flies had periods longer than 24 h. Therefore, the mean period values of the respective genotypes were considered to calculate the circadian time (CT) for dissections to detect PER oscillations in the LNv. For quantifying PER oscillations in DD, LD-reared flies were dissected at CT23-24 (CT23) and CT11-12 (CT11) at different ages: all the five genotypes at 3d and *pdf>Q0,HSP40* and *pdf>Q128,HSP40* also at 9d. These samples were co-stained with anti-PDF to aid in identifying LNv and anti-HTT. The sample sizes were determined empirically (Table S3). There was no blinding during sample preparation.

### Image acquisition and analysis

The number of PDF^+^ and PER^+^ LNv and the form of expHTT in the LNv were noted on manual observation of the samples using a Zeiss Axio Observer Z1 epifluorescence microscope with the 63X/oil 1.4NA objective and without blinding. The collected data were then cross-verified with images captured using a 40X/oil 1.3NA objective as a *z*-stack of 1µm interval. The PDF stained sLNv and lLNv were distinguished based on their anatomical location, size, and staining pattern. For quantification of PER intensity and expHTT inclusions, the lamp intensity and exposure time were kept constant across samples for an experiment. The PER intensity was calculated from the obtained images as described previously in a single-blind manner (Prakash et al., 2017). NIH imaging software ImageJ was used for image analysis and quantification (Schneider et al., 2012). For representative images, confocal *z*-stacks were captured using Zeiss LSM 880, keeping the PMT gain, offset and laser power below saturating pixels. In the representative images, brightness/contrast adjustments have been applied to the whole image for better visualisation of the LNv, especially the sLNv, which show less intense and fainter PDF staining than the lLNv.

### Categorisation and quantification of expHTT forms

The number of sLNv and lLNv with different forms of expHTT was noted in each hemisphere. The expHTT forms appearing in each LNv soma were classified based on their appearance under the light microscope as follows: uniform diffuse expHTT staining (Diff), expHTT appearing as puncta-like shiny specks of varying sizes was referred to as inclusions (Inc), diffuse expHTT with a few puncta-like inclusions (Diff+Inc) and without expHTT staining (No HTT) (Fig. S1A top-panels). With overexpression of HSP40, we also observed a hitherto unreported form of expHTT, oval with a compact appearance, henceforth referred to as expHTT spot or spot-like expHTT (Spot) (Fig. S1A second-row). Also observed less frequently was an LNv with an expHTT Spot and the canonical inclusions, giving the Spot a distorted appearance. Hence, such forms of expHTT were included under the Inc category. If the spot appeared amidst a diffused expHTT distribution, mostly seen in lLNv of young flies, it was designated as Diff+Spot (Fig. S1A second-row). Two sets of information can be gathered from such an exercise of labelling expHTT types per LNv. One is at the level of cells, wherein within each hemisphere, the proportion of sLNvs or lLNvs with different forms of expHTT was noted. Such a comparison (Intra-hemisphere) gives an idea of the within-hemisphere variation in LNv expHTT distribution. Our experimental replicates are at the level of hemispheres and not cells; so, the within-hemisphere expHTT diversity in cells is only qualitative information. The second set of information is at the level of hemispheres. Each hemisphere was allotted a particular category depending on the most predominant expHTT form found sLNv (or lLNv) (Fig. S1B). The hemisphere categorisation (inter-hemisphere) based on predominant expHTT form in LNv (sLNv or lLNv) was as follows: predominantly diffuse distribution (Fig.S1B top-left), an equal distribution of diffused and inclusions (Diff+Inc) (Fig. S1B top, middle and right), predominantly having Diff+Spot (Fig. S1B second-row, left), predominantly had spots (Spot) (Fig. S1B second-row, middle and right), a near equal mix of diffuse, inclusions and spots (Diff+Spot+Inc) (Fig. S1b third row, left), an equal distribution of spot and inclusions (Spot+Inc) (Fig. S1b third-row, middle and right), or predominantly had inclusions (Inc) (Fig. S1b, fourth-row, middle and right). Hemisphere-level categorisation just described is at the level of experimental replicates. It, therefore, makes room for quantitative statistical analysis and allows for comparisons between genotypes and among ages of the relative proportions of hemispheres enriched in one form of expHTT in sLNv or lLNv versus other forms of expHTT. Upon such a hemisphere-level categorisation, hemispheres in which sLNv or lLNv were without expHTT never dominated, so the NoHTT category does not exist. The arrangement order used in the figures describing various expHTT forms is based on observations made post-hoc as to the appearance and predominance of various expHTT forms in LNv over time (Fig. S1C). For example, in most *pdf>Q128* samples, sLNv show Diff expHTT as larvae, and young adults mostly exhibit Diff+Inc in lLNv, followed by Inc as they age. For *pdf>Q128,HSP40* flies mostly show expHTT as Diff in young flies. With age, different combinations of Diff, Spot and some Inc appear and dominate (mostly non-Inc forms), and exclusively Inc becomes more prominent only in much older flies.

### Quantification of expHTT inclusions

The expHTT number and size of expHTT inclusions were quantified using ImageJ. Maximum intensity projection images were converted to 8-bit images. Their backgrounds were subtracted (rolling ball radius of 10 pixels), unsharp mask filter applied (radius of 1 pixel, mask weight of 0.7), further processed to sharpen the image, find edges, and then the intermodes-threshold was applied. The area in and around the LNv was then marked. The analyse particles tool with size specification of 1 to infinity (in µ) was used to obtain measures of inclusion number and the size of inclusions for each hemisphere. A lower limit of 1µ was set for size to avoid false positives and background specks. The quantification method did not distinguish between expHTT Inc and Spots, resulting in Spots being included in the inclusion number and size quantifications.

### Statistical analysis of cellular features

For comparing PDF^+^ or PER^+^ LNv numbers between genotypes or between ages, the Kruskal-Wallis test of ranks, followed by multiple comparisons of mean ranks, was used. The change in the shape of frequency distributions of PDF^+^ and PER^+^ LNv numbers between genotypes was assessed using Kolmogorov-Smirnov tests with α = 0.05, followed by Bonferroni correction on pair-wise *p* values. For comparing inclusion numbers between genotypes or across age, multi-factor ANOVA followed by Tukey’s HSD post-hoc test were used on transformed data. Since the variances were not homogenous for inclusion size comparisons, the transformed data sets, primarily normal, were subjected to Welch’s ANOVAs followed by Games-Howell post-hoc tests (McDonald, 2014). The data sets were either transformed (where required) to analyse the status of PER oscillations between the time-points CT23 and CT11 for a genotype or the PER intensities at a time-point between genotypes or directly analysed using one-way ANOVA, followed by Tukey’s HSD post-hoc test wherever necessary. To compare the proportion of hemispheres predominated by different expHTT forms between genotypes for an age or between ages for a genotype, m x n Fisher’s Exact tests were used. Following this, wherever necessary, multiple specific 2×2

Fisher’s Exact test sets with Benjamini Hochberg procedure on all relevant comparisons were applied (using R). The proportion of hemispheres with Spot^+^ LNv between ages was compared using multiple 2×2 Fisher’s exact tests followed by Bonferroni corrections. Spot sizes between ages for sLNv or lLNv were analysed by one-way ANOVA (on transformed data, if required), followed by Tukey’s HSD tests. For Spot sizes between sLNv and lLNv, a factorial ANOVA with LNv and age as fixed factors were carried out on transformed data, followed by Tukey’s HSD.

OriginPro 8 (Origin(Pro)), Sigma Plots 11.0 (SigmaPlot), and Adobe InDesign 3.0 (AdobeIndesignCS) were used for making figures.

## Results

### Overexpression of HSP40 or HSP70 delays arrhythmicity in flies expressing expHTT in LNv

Flies expressing expHTT in LNv are arrhythmic immediately upon entering DD25 (Fig. 1A second-row, left). Overexpressing heat shock proteins HSP40 or HSP70 in *pdf>Q128* flies delayed the onset of this arrhythmicity (Fig. 1A second-row, middle and right). Most flies co-expressing HTT-Q128 with HSP40 were rhythmic during the early age window (henceforth, AW1) and mid-age window (henceforth, AW2), their mean rhythmicities being comparable to controls expressing HTT-Q0 and significantly higher than *pdf>Q128* (Fig. 1B). However, their rhythmicity during a later age window (henceforth AW3) declined significantly compared to controls and earlier AWs and was like *pdf>Q128*. Despite a drop in rhythmicity compared to AW1, most *pdf>Q128,HSP40* flies remained rhythmic during AW2. *pdf>Q128,HSP70,* had significantly higher rhythmicity than *pdf>Q128* in AW1 and AW2 (Fig. 1B). However, it was like controls only in AW1, beyond which it declined. Notably, in AW2, while about 50% of *pdf>Q128,HSP70* remained rhythmic, their rhythmicity was significantly lower than *pdf>Q128,HSP40*. In AW3, like *pdf>Q128,HSP40* and *pdf>Q128, pdf>Q128,HSP70* also had poor rhythmicity. While the rhythmicity of *pdf>Q128,HSP40* was like background controls across AW1 and AW2 (Figs S2A, middle-column, 1C, left), that of *pdf>Q128,HSP70* was control-like only during AW1 (Figs S2A, right-column, 1D, left). Unlike the sharp fall in rhythmicity of *pdf>Q128,HSP70* in AW2, *pdf>Q128,HSP40*, despite a progressive reduction in rhythmicity with age, showed a significant fall in mean rhythmicity only in AW3 (Fig. 1A). Thus, the rescue of rhythmicity by HSP40 lasted longer than HSP70.

**Fig. 1.**
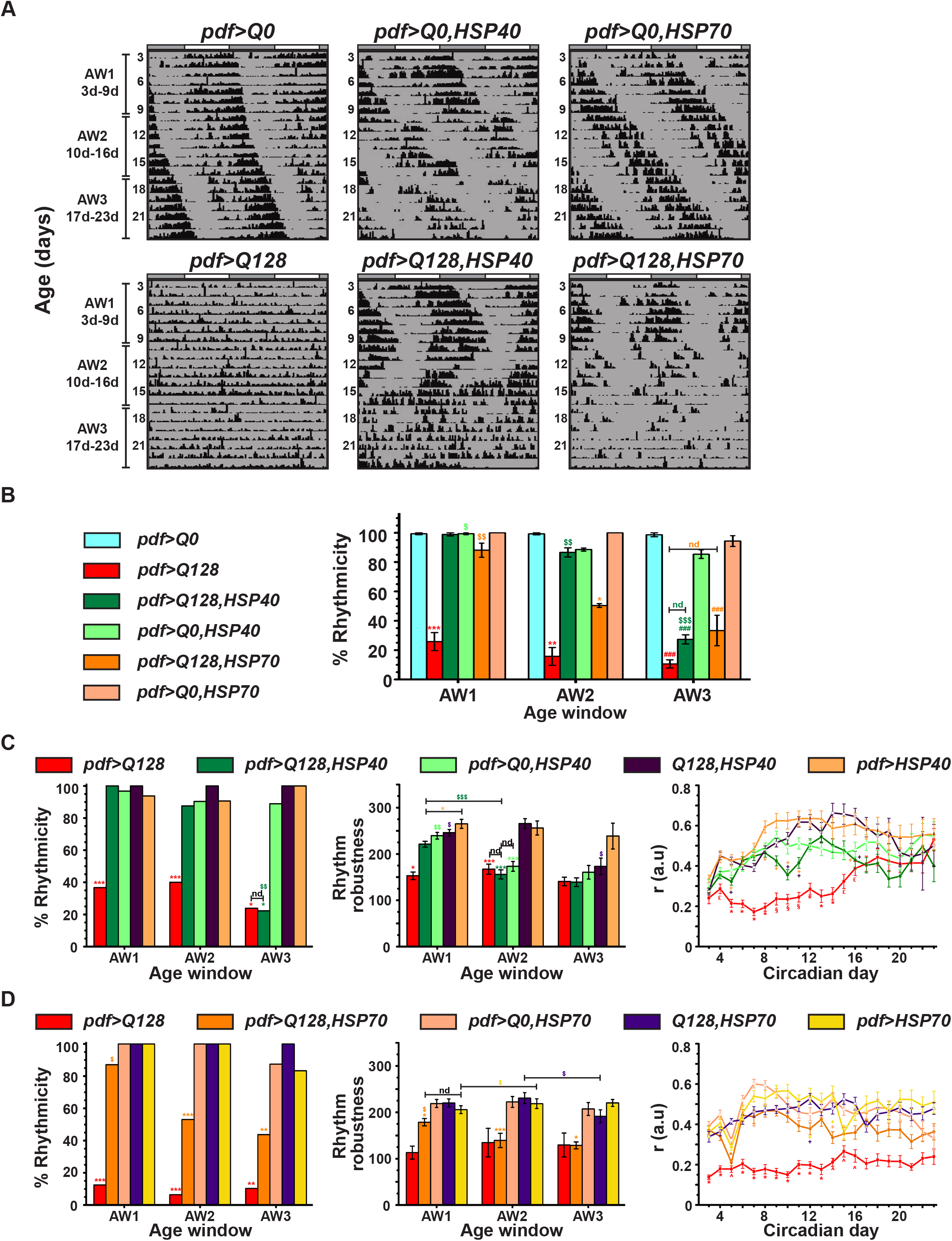
In *pdf>Q128* flies, HSP40 overexpression leads to sustained behavioural rhythms, while HSP70 overexpression leads to early-age rhythmicity. (A) Representative double- plotted actograms for flies showing activity data for 21d (age 3d-23d) in DD at 25°C for *pdf>Q128*, *pdf>Q128,HSP40*, *pdf>Q128,HSP70* and their respective *Q0* controls. Data of 21d is divided into three 7d age windows (AWs), shown on the left. The white and grey bars above actograms indicate the light and dark phases of prior LD. (B) The percentage rhythmicity averaged over at least three independent runs plotted across AWs. (C) and (D) Comparison of percentage rhythmicity (left), mean rhythm robustness (middle), and mean ‘*r*’ value (right) between genotypes across age with the main experimental genotype being *pdf>Q128,HSP40* (C) and *pdf>Q128,HSP70* (D). *pdf>Q128* has not been considered for between-genotype statistical comparisons of robustness during AW3 (C) and across age (D), as very few flies were rhythmic. Across all panels coloured symbols represents statistically significant differences: coloured * indicates age-matched differences of the respective-coloured genotype from all other genotypes or indicated genotype, coloured # indicates age-matched differences of the respective-coloured genotype from all *Q0*-containing controls and coloured $ indicates differences across age for the respective-coloured genotype. Statistical significance at single- symbol *p*<0.05, dual-symbol *p*<0.01 and triple-symbol at *p*<0.001. nd indicates not different. For the bottom-panels of ‘*r*’ values, red-coloured symbols indicate significant differences at *p* <0.05 of *pdf>Q128* from * all other genotypes, § from all genotypes except *pdf>Q128,HSP40*, ˄ from all genotypes except *pdf>Q128,HSP70*, £ from all non-expanded controls and * (orange) of *pdf>Q128,HSP70* from all other genotypes. Coloured + near the error bar of a data-point indicates significant differences at *p* <0.05 of the respective-coloured genotype from the data-point genotype. Error bars are s.e.m.

In AW1, the rhythmic flies of *pdf>Q128,HSP40* had robust rhythms comparable to most controls and significantly higher robustness than *pdf>Q128* (Fig. 1C, middle). In AW2, the robustness of rhythmicity of *pdf>Q128,HSP40* dropped lower than both parental controls and comparable to *pdf>Q128*. Overall, the overexpression of HSP40 in *pdf>Q128* flies rescued both rhythmicity and rhythm robustness in AW1. In AW1, the rhythmic *pdf>Q128,HSP40* and *Q128,HSP40* flies had longer periods than other controls (Figs 1A, S2A, B). The *pdf>Q128,HSP40* flies had a significantly better activity consolidation than *pdf>Q128* flies across ages 5d-8d and 11-14d (Fig. 1C, right). However, this improved consolidation was comparable to controls at 6-7d and 12-15d. Thus, overexpression of HSP40 in *pdf>Q128* flies rescues both rhythmicity and rhythm strength at an early age. Activity rhythms persist until the middle-age, postponing arrhythmicity onset by more than two weeks. These results indicate that HSP40 is a potent suppressor of expHTT-induced circadian behavioural arrhythmicity.

The rhythmic flies of *pdf>Q128,HSP70* exhibited weak rhythms than controls across AWs and robustness in older AWs was lower than AW1 (Fig. 1D, middle). Controls *pdf>Q0,HSP70* and *pdf>HSP70* mostly had longer periods than rhythmic *pdf>Q128,HSP70* and *Q128,HSP70* across AWs (Figs 1A, S2A, C). The activity consolidation of *pdf>Q128,HSP70* was significantly better than *pdf>Q128* across 4d-5d, 6d-13d and comparable to controls at most of these ages (Fig. 1D, right). Thus, the overexpression of HSP70 in *pdf>Q128* flies improves their early-age rhythmicity and activity consolidation. In contrast to the rescue with HSP40 overexpression, rhythm rescue with HSP70 overexpression at an early age did not extend to the middle-aged. Hence, HSP70 is a partial suppressor of expHTT-induced circadian behavioural arrhythmicity and is less efficient than HSP40.

### Co-expression of HSP40 and HSP70 synergistically improves consolidation of activity rhythms in flies expressing expHTT in LNv

Previous studies show that co-expression of HSP40 and HSP70 has a synergistic effect of providing a greater effect on neurodegenerative features than expressing HSP40 or HSP70 individually (Chan et al., 2000; Jana, 2000; Kobayashi et al., 2000; Muchowski et al., 2000; Sittler et al., 2001; Bailey et al., 2002; Bonini, 2002; Rujano et al., 2007). Therefore, we asked whether overexpression of both HSPs provides a greater rescue (e.g. sustained robust rhythms across AWs) than expressing each alone. In AW1, flies expressing both HSPs in the presence of expHTT, i.e. *pdf>Q128,HSP40,70* were mostly rhythmic, comparable to *pdf>Q128,HSP40* and *pdf>Q128,HSP70* and significantly better than *pdf>Q128* (Fig 2A, 2B, left). However, in AW2, the percentage rhythmicity of *pdf>Q128,HSP40,70*, like that of *pdf>Q128,HSP70*, dropped significantly compared to AW1, while remaining higher than *pdf>Q128*, but lower than *pdf>Q128,HSP40* and *pdf>Q0,HSP40,70* (Fig 2B, left). In AW3, the rhythmicity percentage of *pdf>Q128,HSP40,70* declined further and like the single HSP overexpression, comparable to *pdf>Q128*. In AW1, the rhythmic *pdf>Q128,HSP40,70* flies exhibited robust rhythms comparable to control *pdf>Q0,HSP40,70* and the single rescue *pdf>Q128,HSP40* (Fig 2B, middle). Its period was longer than other genotypes (Figs 2A, S2D). *pdf>Q128,HSP40,70* and *pdf>Q128,HSP70* had weaker rhythms than *pdf>Q128,HSP40* in AW1 and from *pdf>Q0,HSP40,70* also in AW2 (Fig 2B, middle). Interestingly, the extent of activity consolidation ‘*r*’ of *pdf>Q128,HSP40,70* was significantly greater than *pdf>Q128* and both *pdf>Q128,HSP40* and *pdf>Q128,HSP70* during most of the early age (Fig 2B, right). ‘*r*’ of *pdf>Q128,HSP40,70* remained higher than *pdf>Q128* and comparable to *pdf>Q0,HSP40,70* across age. Thus, although overexpression of both HSP40 and HSP70 in *pdf>Q128* did not have a synergistic effect on percentage rhythmicity *per se*, there was a synergistic improvement in early-age activity consolidation.

**Fig. 2.**
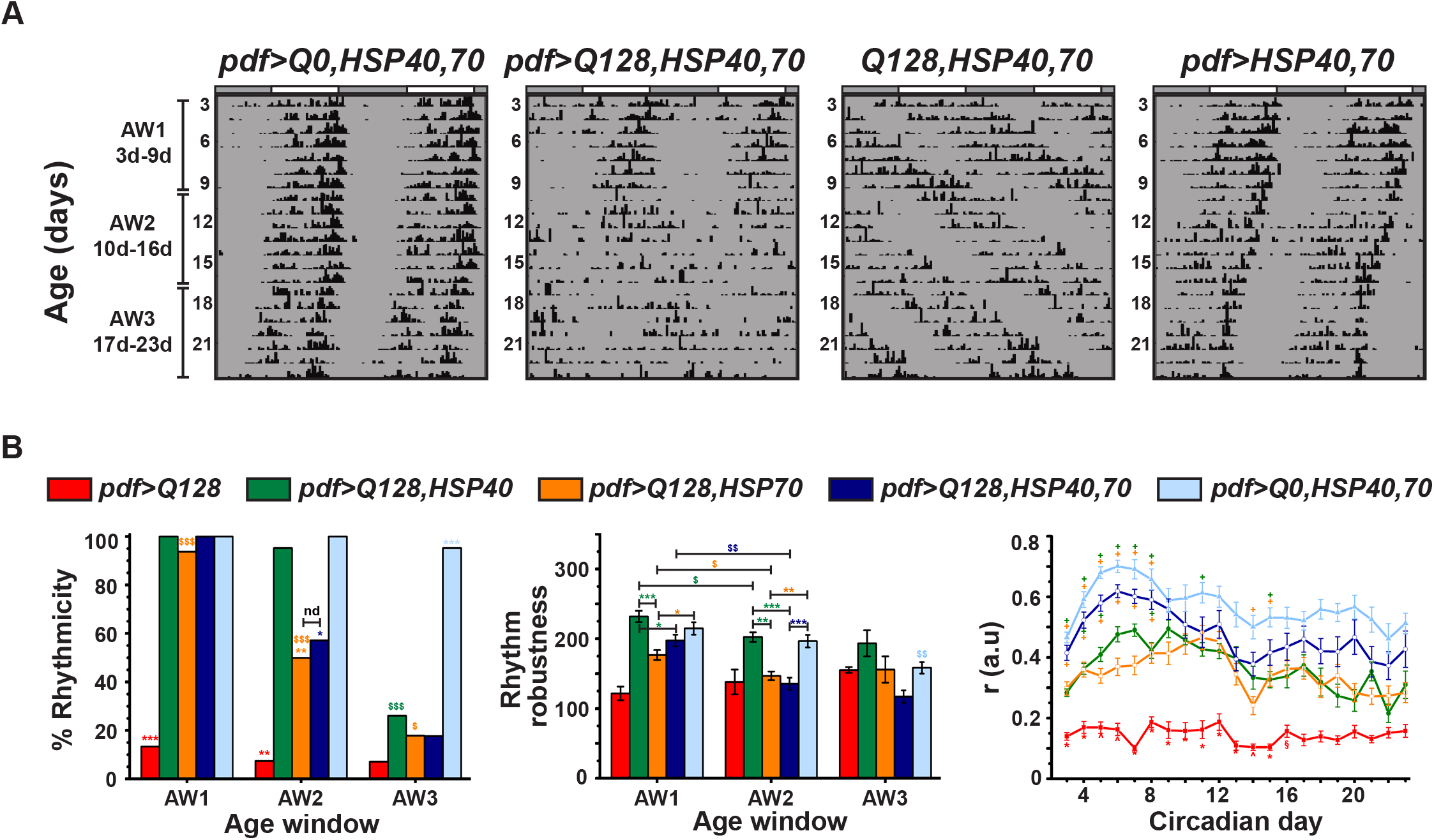
*pdf>Q128* flies co-expressing both HSP40 and HSP70 have greater daily activity consolidation than either HSP expressed alone. (A) Representative double-plotted actograms for flies showing activity data for 21d (age 3d-23d) in DD at 25°C for *pdf>Q128,HSP40,70* and its controls. (B) The percentage of rhythmic flies (left), mean rhythm robustness (middle) and mean ‘*r*’ value (right) comparing genotypes across age. *pdf>Q128* and AW3 are omitted from between-genotype statistical tests for robustness. Across all panels coloured symbols represents statistically significant differences: coloured * indicates age- matched differences of the respective-coloured genotype from all other genotypes or indicated genotype, coloured # indicates age-matched differences of the respective-coloured genotype from all *Q0*-containing controls and coloured $ indicates differences across age for the respective-coloured genotype. Statistical significance is at single-symbol *p*<0.05, dual-symbol *p*<0.01 and triple-symbol at *p*<0.001. nd indicates not different. For the bottom-panels of ‘*r*’ values, red-coloured symbols indicate significant differences at *p* <0.05 of *pdf>Q128* from * all other genotypes, § from all genotypes except *pdf>Q128,HSP40*, ˄ from all genotypes except *pdf>Q128,HSP70*, and * (orange) of *pdf>Q128,HSP70* from all other genotypes. Coloured + near the error bar of a data-point indicates significant differences at *p* <0.05 of the respective- coloured genotype from the data-point genotype. nd is not different. Error bars are s.e.m.

### HSP40 overexpression in flies expressing expHTT in LNv rescues PDF^+^ sLNv soma numbers

We then investigated whether overexpression of HSP40 or HSP70 in *pdf>Q128* also rescues LNv cellular features. As described previously (Sheeba et al., 2010; Prakash et al., 2017) and as is also shown here, *pdf>Q128* flies had a loss of PDF from the sLNv soma from an early age, while PDF in lLNv soma was unaltered (Figs 3, top, 4A). In contrast, flies overexpressing HSP40, the *pdf>Q128,HSP40* showed significantly higher PDF^+^ sLNv soma numbers than *pdf>Q128* and indistinguishable from control *pdf>Q0,HSP40* across age (Figs 3, middle panel-sets, 4A, left). The shapes of the frequency distributions of PDF^+^ sLNv numbers for *pdf>Q128,HSP40* across age were left-skewed, like controls, with most hemispheres having 4-5 sLNv and differed significantly from the right-skewed distribution of *pdf>Q128* (Fig 4B). In contrast, at 3d and 9d, the PDF^+^ sLNv soma numbers of *pdf>Q128,HSP70* were diminished like that of *pdf>Q128* and significantly lower than control *pdf>Q0,HSP70* and from *pdf>Q128,HSP40* (Figs 3, bottom panel-sets, 4A, left). Mirroring the mean PDF^+^ sLNv soma numbers was the shape of their distributions at both ages: *pdf>Q128,HSP70*, was like *pdf>Q128* and different from controls (Fig 4C). The PDF^+^ lLNv soma numbers were comparable for all the genotypes across age (Figs 3, arrow-heads, 4A, right). Thus, overexpression of HSP40, but not HSP70, completely rescues PDF^+^ sLNv numbers. The sustained circadian rhythm rescue in *pdf>Q128,HSP40* accompanied by the rescue of the circadian output neuropeptide PDF in the soma of pacemaker neurons sLNv, suggests HSP40 as a disease-modifier effective in restoring cellular function as well as its associated behaviour. The lack of rescue of PDF^+^ sLNv soma in *pdf>Q128,HSP70* at an early age despite the persistence of circadian activity rhythms suggests an unconventional mode of rhythm restoration by HSP70 in the absence of somal PDF in the sLNv.

**Fig. 3.**
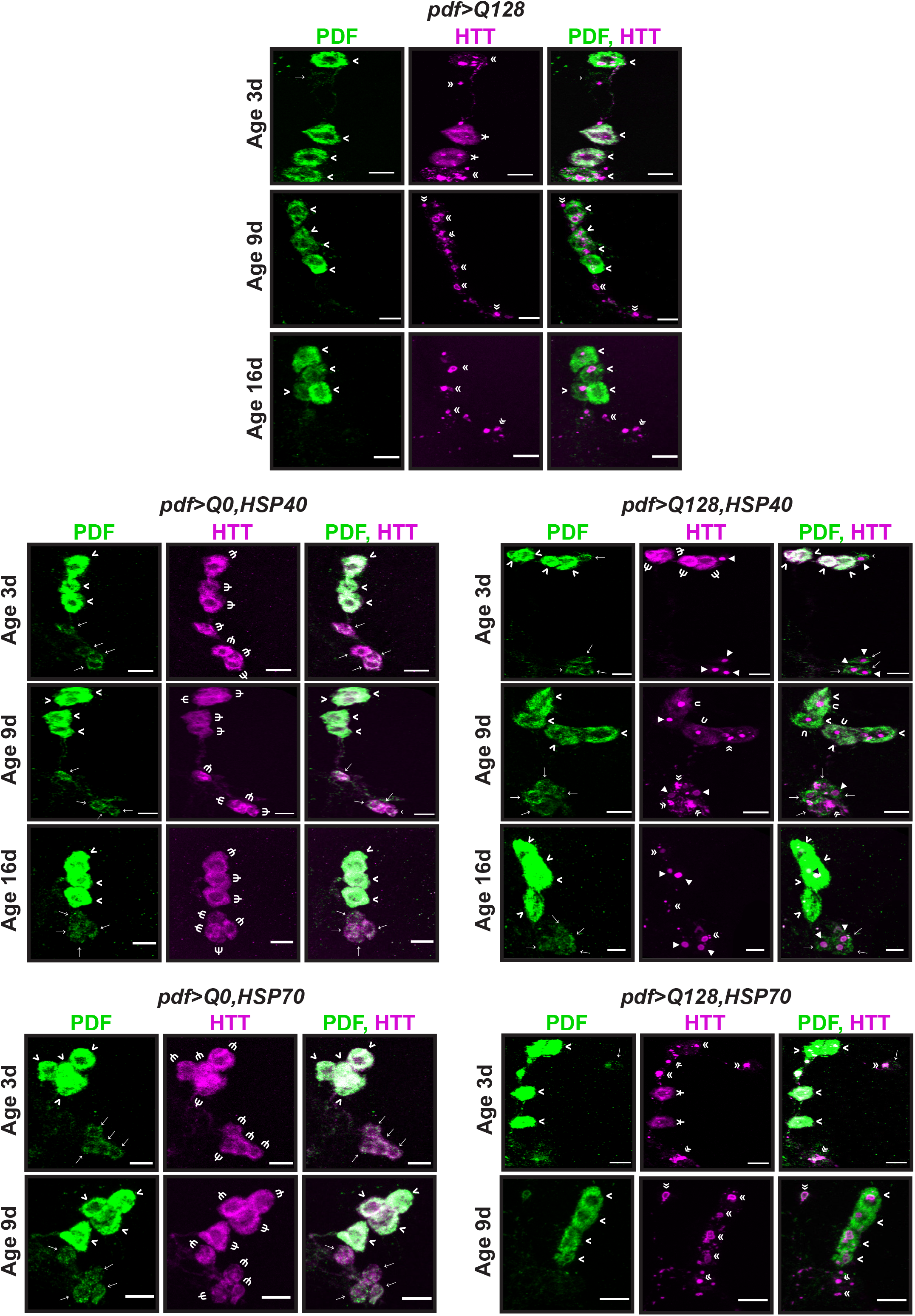
*pdf>Q128* flies overexpressing HSP40 retain PDF^+^ sLNv soma across age. Representative images of adult fly brains stained for PDF (green) and HTT (magenta) in LNv at 3d, 9d and 16d for *pdf>Q128* (top panel-sets), *pdf>Q0,HSP40* (middle-left panel-sets), *pdf>Q128,HSP40* (middle-right panel-sets), and at 3d and 9d for *pdf>Q0,HSP70* (bottom-left panel-sets) and *pdf>Q128,HSP70* (bottom-right panel-sets). Indicated in the images are sLNv soma (→ arrows), lLNv soma (> arrowheads), diffuse expHTT (Ψ psi), diffuse+inclusions expHTT (Ұ), diffuse+spot expHTT (υ upsilon), spot expHTT (◄ triangles) and expHTT inclusions (« double arrowheads) for the five genotypes. Scale bars are 10 µm.

**Fig. 4.**
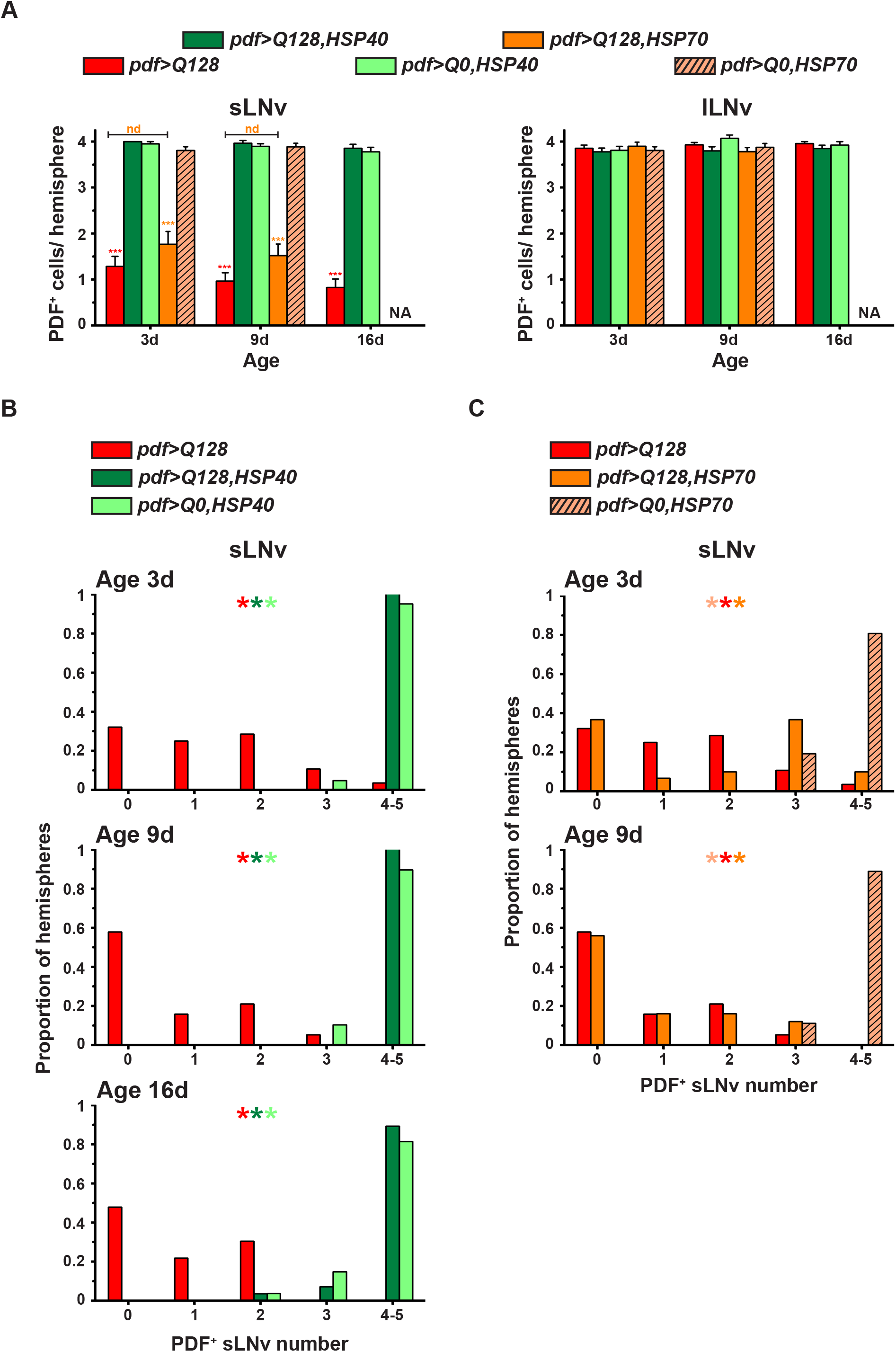
*pdf>Q128* flies overexpressing HSP40 have control-like PDF^+^ sLNv soma numbers. (A) Mean number of PDF^+^ sLNv soma (left) and lLNv soma (right) across three ages. Symbols indicate significant differences, * for age-matched, inter-genotype differences and $ for differences between ages for each genotype: * (red) of *pdf>Q128* from all other genotypes except *pdf>Q128,HSP70* and * (orange) of *pdf>Q128,HSP70* from other genotypes except *pdf>Q128* at * *p*<0.05, ** *p*<0.01, *** *p*<0.001. NA is not applicable; since early-age-rescue of PDF^+^ LNv was not seen in *pdf>Q128,HSP70*, 16d dissections were not done for it. (B) and (C) Frequency distribution of the proportion of hemispheres with 0 to 5 PDF^+^ sLNv soma numbers comparing *pdf>Q128* with *pdf>Q128,HSP40* and *pdf>Q0,HSP40* at 3d, 9d and 16d (B) and with *pdf>Q128,HSP70* and *pdf>Q0,HSP70* at 3d and 9d (C). Coloured multiple * indicates significantly different distribution shapes between genotypes, with the first colour of the reference genotype and the subsequent colours of genotypes differing from the reference at *p*<0.01. nd indicates not different. Error bars are s.e.m.

### HSP40 overexpression in *pdf>Q128* flies reduces the inclusion form of expHTT in favour of a new form

HSPs are molecular chaperones and are known to interfere with various stages of aggregation and modify the nature, conformation, and solubility of expHTT inclusions (rral et al., 2004; Wyttenbach and Arrigo, 2009; Lotz et al., 2010; Arrasate and Finkbeiner, 2012). We used immunocytochemistry and light microscopy to determine whether HSP overexpression modifies the expHTT forms detected in the LNv of *pdf>Q128*. As detailed in the methods section, the expHTT forms present in each sLNv or lLNv were categorised based on their appearance. Interestingly, *pdf>Q128* with overexpressed HSP40 had an additional expHTT form that has not been observed in these flies and instances of which seem unreported in the literature. Visually and qualitatively, this form of HTT-Q128 appeared as a compact oval and was excluded from the cytoplasmic PDF staining (Figs 3, middle row, right panel-sets, see triangles, 5A, B). We refer to this as the “Spot” form of expHTT. PER at CT23 is mainly nuclear, and this compact expHTT Spot form appears in the vicinity of nuclear PER and might be peri-nuclear (Fig. 5 A, B). The Spot form was also restricted to a single structure per LNv.

**Fig. 5.**
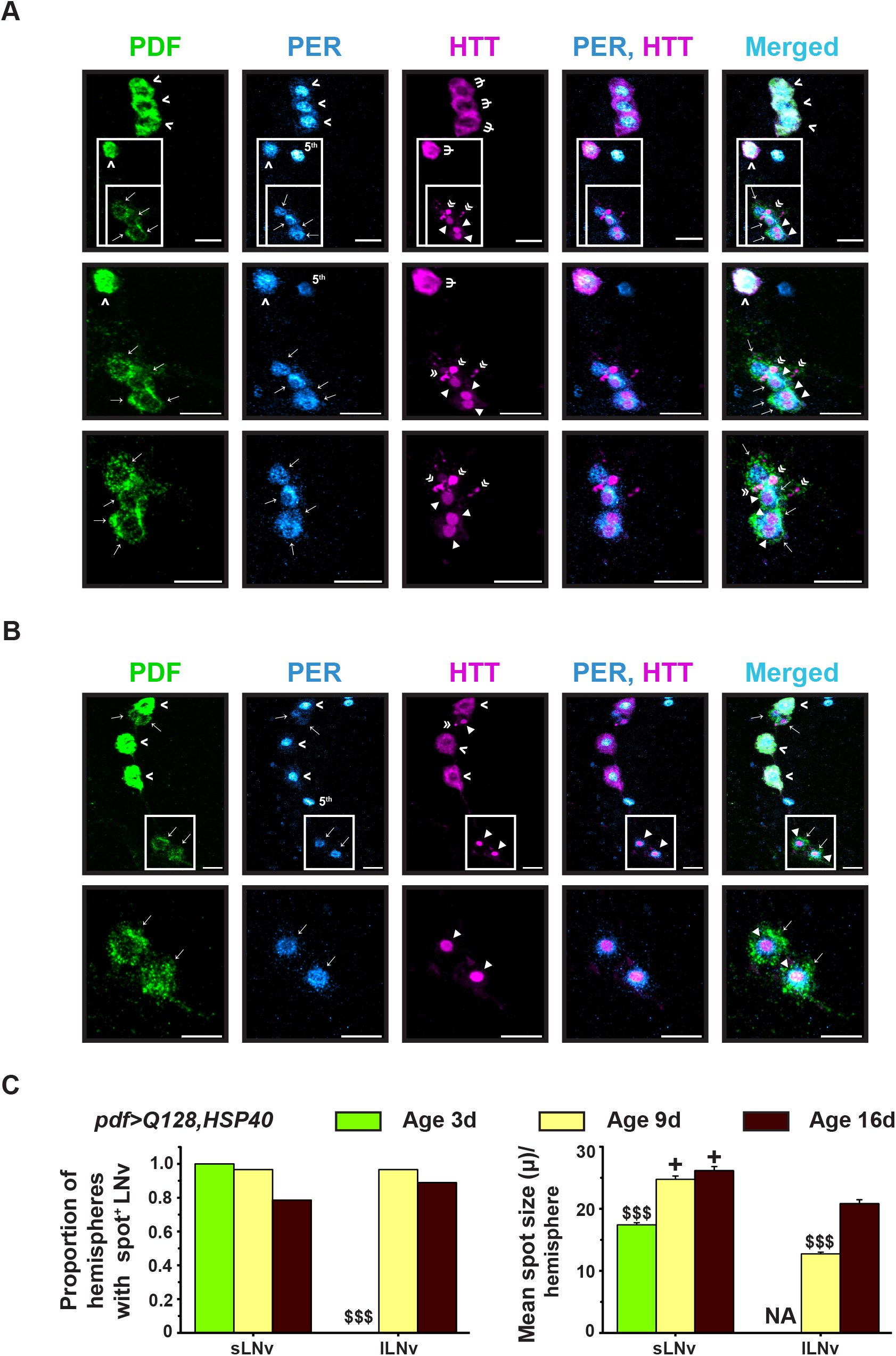
*pdf>Q128* flies overexpressing HSP40 show the presence of a novel expHTT form, the Spot. (A) and (B) Two sets of representative images of 3d adult brains of *pdf>Q128,HSP40* stained for PDF (green), PER (cyan hot) and expHTT (magenta) showing better resolved expHTT spots, where marked rectangles in each panel are enlarged in the subsequent panels below. Indicated in the images are sLNv soma (→ arrows), lLNv soma (> arrow-heads), diffuse expHTT (Ψ psi), spot expHTT (◄ triangles), expHTT inclusions (« double arrow-heads) and the PDF^-^, PER^+^ 5^th^ sLNv. (C) Across three ages, the proportion of hemispheres with spots in sLNv or lLNv (left) and the mean spot sizes in sLNv and lLNv (right) are compared. $ depicts the difference of one age from other ages at $$$ *p*<0.001 and + indicates age-matched differences between sLNv and lLNv at *p*<0.0001. NA is not applicable. Error bars are s.e.m. Scale bars are 10 µm.

Only the *pdf>Q128,HSP40* showed the presence of expHTT Spots (Fig. 3). Within each hemisphere, Spot expHTT was present in ∼75% sLNv (3 of 4) at 3d and ∼50% sLNv (2 of 4) at 9d and 16d (Fig. S3A, top). At 3d and 9d, nearly every hemisphere of *pdf>Q128, HSP40* had at least one sLNv with Spot expHTT, which decreased to ∼ 75% at 16d (Fig. 5C, left). In lLNv of *pdf>Q128,HSP40*, expHTT Spot was absent at 3d, detected in ∼60% lLNv per hemisphere (2-3 of 4) at 9d as Diff+Spot and in ∼50% lLNv at 16d as a distinct Spot (Fig. S3A, bottom). Across samples of *pdf>Q128,HSP40*, most of the hemispheres showed the presence of Spot in at least one lLNv (as Diff+Spot or Spot) at 9d and 16d (Fig. 5C, left). On average, sLNv of older flies had significantly bigger spots (∼25 µ) than 3d flies (∼17 µ), and in lLNv, a similar trend was seen, with larger spots at 16d (∼20µ) than 9d (∼12µ) (Fig. 5C right). The spots in the sLNv were larger than those in age-matched lLNv (Fig 5C, right).

Comparing the between-hemisphere distribution of predominant expHTT forms in LNv, it is apparent that at 3d and 9d, inclusions dominated in both the sLNvs and lLNvs of *pdf>Q128* and *pdf>Q128,HSP70* (Figs 3, 6A, B). In contrast, in the LNv of *pdf>Q128,HSP40*, non-inclusion forms of expHTT like Diff and Spot dominated over exclusively-Inc (Fig. 6A, B). At both ages, the overall distributions of expHTT forms in sLNv and lLNv of *pdf>Q128,HSP40* differed significantly from those of *pdf>Q128* and *pdf>Q128,HSP70* (Fig. 6A). We then compared the relative proportion of hemispheres in various pair-wise-category combinations. Specifically, at 3d, the proportion of hemispheres dominated by Inc expHTT in sLNv relative to forms was significantly higher in *pdf>Q128* and *pdf>Q128,HSP70* than *pdf>Q128,HSP40*, which mostly had hemispheres dominated by Spot expHTT and to a lesser extent Spot+Inc in the sLNv (Fig. S3B, top). At 9d, nearly 50% of *pdf>Q128,HSP40* hemispheres still had Spot-predominant sLNv either as Spot or Spot+Inc expHTT, while a similar proportion of hemispheres also had Inc enriched sLNv (Fig. S3B, bottom). It is of note that, by 9d, more than 50% of *pdf>Q128* and *pdf>Q128,HSP70* had no PDF^+^ sLNv, with mean number ∼1, whereas nearly all *pdf>Q128,HSP40* had 4 PDF^+^ sLNv (Figs. S3B- indicated at the bottom of each bar are the mean PDF^+^ LNv numbers).

**Fig. 6.**
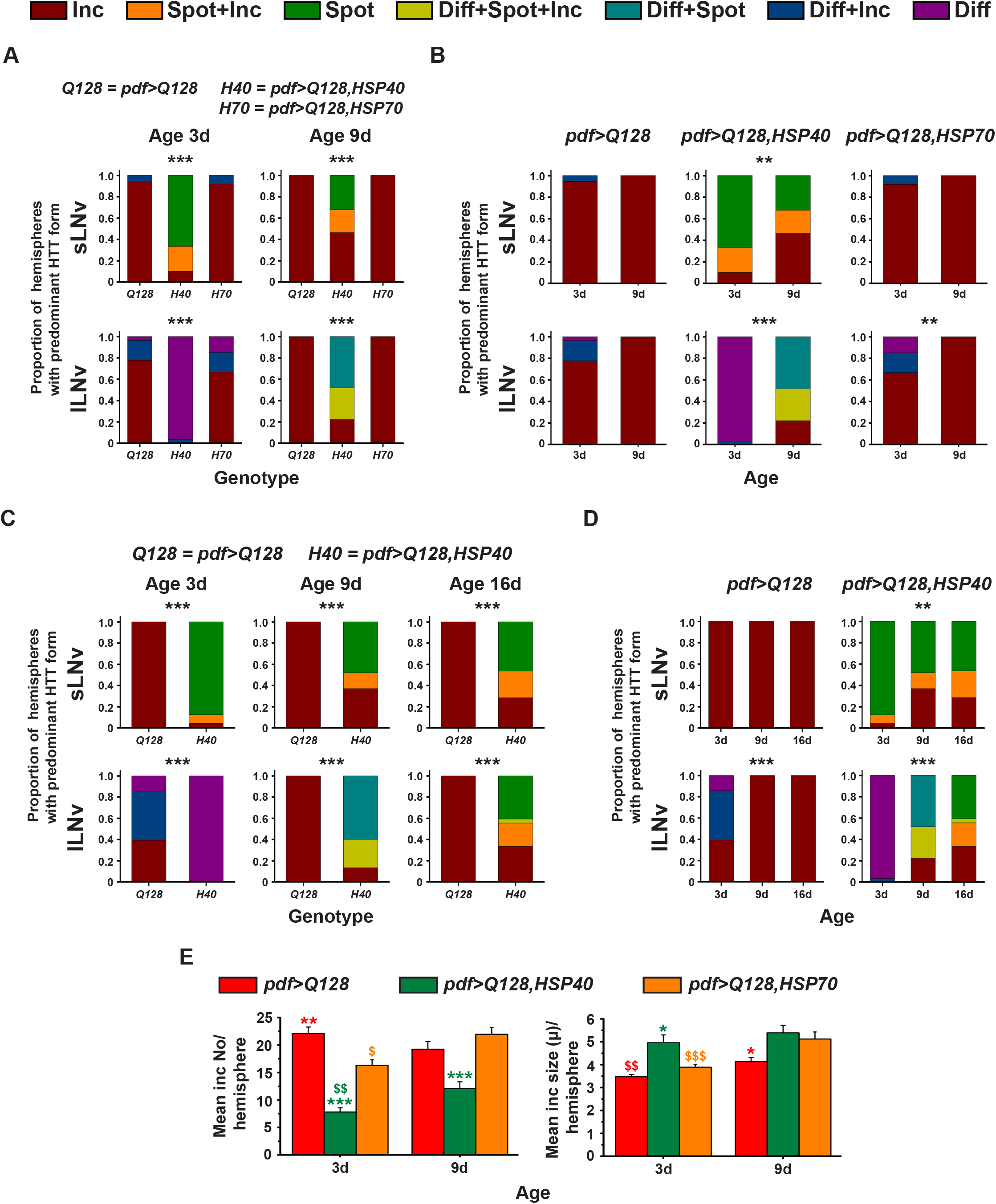
*pdf>Q128* flies overexpressing HSP40 have fewer hemispheres with expHTT inclusion enriched LNv and reduced expHTT inclusions numbers. (A) and (B) The proportion of hemispheres dominated by different expHTT forms in sLNv (top) or lLNv (bottom) is plotted on the *y*-axis to describe the between-hemispheres (inter-hemisphere) distribution of predominant expHTT forms. This proportion is plotted at 3d and 9d against three genotypes (A) or for each genotype against ages 3d and 9d (B). * indicates significant changes in relative distributions of expHTT forms between genotypes (A) or between ages (B) at ** *p*<0.01 and *** *p*<0.0001. The relevant pair-wise comparisons of (A) are plotted in Fig. S3B, C. (C) and (D) are similar to (A) and (B), comparing the three ages, 3d, 9d, and 16d for *pdf>Q128* and *pdf>Q128,HSP40*. The relevant pair-wise comparisons of (D) are plotted in Fig. S4 A-C. (E) Comparison of mean inclusion number per hemisphere (left) and mean inclusion size per hemisphere (right) for three genotypes at 3d and 9d. Coloured * indicates statistically significant age-matched differences between genotypes: * (olive-green) from *pdf>Q128,HSP40* and * (red) from *pdf>Q128*, coloured $ represent differences across age for the respective-coloured genotype. Statistical significance is at single-symbol *p*<0.05, dual- symbol *p*<0.01 and triple-symbol *p*<0.001. Error bars are s.e.m.

The proportion of hemispheres with Inc predominant lLNv relative to other forms of expHTT was significantly higher in *pdf>Q128* and *pdf>Q128,HSP70* than *pdf>Q128,HSP40* at both 3d and 9d (Fig. S3C, top-left and bottom). *pdf>Q128,HSP40* at these ages favoured mostly non-Inc form enriched lLNv. At 3d, most of the hemispheres of *pdf>Q128,HSP40* had Diff expHTT in lLNv, and by 9d, Spot expHTT appeared, giving rise to hemispheres predominated by mostly non-Inc expHTT forms in lLNv, namely Diff+Spot and Diff+Spot+Inc (Fig. S3C). Thus, the inclusion form of expHTT predominates over other forms in LNv across age in *pdf>Q128* and *pdf>Q128, HSP70*, whereas, in *pdf>Q128,HSP40* diffuse and spot forms are prevalent.

To track the progress of these distinct forms of expHTT over a more extended duration in the presence of HSP40, in a separate experiment with *pdf>Q128* and *pdf>Q128,HSP40*, we quantified cellular phenotypes up to 16d. Like the previous experimental results (Fig. 6A, B), across age, the relative proportions of hemispheres with different expHTT forms in sLNv and lLNv differed significantly between genotypes (Fig. 6C, D). Most of the *pdf>Q128* hemispheres had Inc-enriched sLNv across age and Inc-enriched lLNv at 9d and 16d; while in *pdf>Q128,HSP40*, Spot expHTT in various combinations dominated the LNv (Fig. 6C, D). In *pdf>Q128*, as the flies aged, there was a significant reduction in the proportion of hemispheres predominating in Diff or Diff+Inc expHTT in lLNv relative to those predominating in Inc expHTT (Fig. S4A). In hemispheres of *pdf>Q128,HSP40*, Spot expHTT enriched sLNv were present across age (Fig. S4B), and with age, the proportion of hemispheres with predominantly Spot expHTT in sLNv relative to Inc expHTT diminished (Fig. S4B, left). *pdf>Q128,HSP40* showed a significant change across age in the relative proportion of hemispheres enriched in Diff expHTT in lLNv to those enriched with other forms of expHTT in lLNv (Fig. S4C, top-row). Post 3d, Diff expHTT steeply declined in lLNv, making way for Diff+Spot, Diff+Spot+Inc and Inc at 9d and Spot, Spot+Inc and Inc at 16d (Fig. S4C, top-row). From 9d to 16d, the relative proportions of hemispheres of Diff+Spot (and Diff+Spot+Inc) enriched lLNv to that of Inc enriched lLNv decreased significantly (Fig. S4C, middle row, first and second panels). Concomitantly, the relative proportions of hemispheres of Spot (and Spot+Inc) enriched lLNv to that of Inc enriched lLNv increased significantly (Fig. S4C, middle-row, third and fourth panels). Thus, the overexpression of HSP40 in *pdf>Q128* flies decreases the expHTT inclusions in LNv, in favour of expHTT spots. HSP70 overexpression, on the other hand, did not decrease expHTT inclusions in LNv.

In summary, HSP40 overexpression improves LNv health by reducing inclusions of expHTT in favour of a new form of expHTT, the “Spot”, and preserving PDF^+^ sLNv. The Spot expHTT might be a relatively less toxic form of expHTT, given the control-like PDF^+^ sLNv numbers of *pdf>Q128,HSP40*. Also, *pdf>Q128,HSP70* and *pdf>Q128* were nearly indistinguishable in the dominance of expHTT inclusions in LNv (Fig 6), suggesting that mechanisms mediating the early-age rhythms upon HSP70 overexpression might not involve mitigation of visible inclusions.

### HSP40 overexpression in *pdf>Q128* flies reduces the number of expHTT inclusions

We quantified the number and size of expHTT inclusions found in and around the LNv. At both 3d and 9d, *pdf>Q128,HSP40* had significantly fewer inclusions than *pdf>Q128* and *pdf>Q128,HSP70* (Fig. 6E, left). At 3d, *pdf>Q128,HSP70* also had fewer inclusions than *pdf>Q128*, but not at 9d. Both *pdf>Q128,HSP40* and *pdf>Q128,HSP70* exhibited an increase in inclusion number with age. Altogether, overexpression of HSP40 or HSP70 in *pdf>Q128* reduces expHTT inclusion numbers, the effect being long-lasting with HSP40.

The mean inclusion size of *pdf>Q128,HSP40* was higher than *pdf>Q128* and *pdf>Q128,HSP70* at 3d, which is likely a reflection of including the relatively large-sized expHTT spots among inclusions during quantification (Fig. 6E, right). Surprisingly, at 9d, *pdf>Q128* had smaller inclusions than *pdf>Q128,HSP40* and *pdf>Q128,HSP70*. Inclusion size of *pdf>Q128* and *pdf>Q128,HSP70* increased with age.

In summary, co-expression of expHTT with HSP40 in the LNv decreases the proportion of hemispheres with inclusion-enriched LNv across age, with a concomitant increase in the proportion of hemispheres enriched in a hitherto unreported Spot form of expHTT in LNv and a decrease in the expHTT inclusion numbers. All the above observations, taken together, will be henceforth referred to as a decrease in the “inclusion load”. Thus, HSP40 overexpression in *pdf>Q128* flies reduces the inclusion load of the LNv.

### HSP40 rescues early-age sLNv PER oscillations in expHTT expressing flies

PER, a central clock protein, is lost from the soma of LNv in *pdf>Q128* flies (Prakash et al., 2017), also seen here, with *pdf>Q128* having significantly fewer PER^+^ sLNv soma at 3d and 9d and almost none at 16d (Figs 7A, left, B, top, S4D, left). We addressed whether the neuroprotective effect of HSP40 overexpression on *pdf>Q128* flies extends to loss of PER and its molecular oscillations in the LNv. *pdf>Q128,HSP40* showed the presence of PER^+^ sLNv, and lLNv soma at 3d and 9d, with control-like numbers (Fig. 7A, left panel-sets, 7B) and their frequency distributions were left-skewed like *pdf>Q0,HSP40* and differed from *pdf>Q128* (Fig. 7C, first and second columns, top and middle rows). However, unlike the rescue of PDF in sLNv soma, PER rescue in the LNv soma was not sustained up to 16d, by which time, *pdf>Q128,HSP40* had significantly fewer PER^+^ sLNv and lLNv and was comparable to *pdf>Q128* (Figs 7B, S4D). The shape of the frequency distribution of PER^+^ sLNv and lLNv soma numbers in *pdf>Q128,HSP40* changes from a control-like left-skew at 9d to a *pdf>Q128*-like shape at 16d (Fig. 7C, first and second columns, middle and bottom rows). *pdf>Q128,HSP70*, did not show rescue of PER^+^ sLNv soma across age. Its mean numbers and frequency distribution were comparable to *pdf>Q128* and significantly differed from *pdf>Q0,HSP70* and *pdf>Q128,HSP40* (Fig. 7A, right panel-sets, B, top, C, right column, top and middle rows). PER^+^ lLNv soma numbers of *pdf>Q128,HSP70* were comparable to *pdf>Q128* at 3d and 9d, control-like at 3d and significantly reduced at 9d (Fig. 7B, bottom). The shape of the PER^+^ lLNv soma distribution of 9d old *pdf>Q128,HSP70*, like that *of pdf>Q128*, differed from the left-skewed distribution of *pdf>Q0,HSP70* (Fig. 7C, bottom-right).

**Fig. 7.**
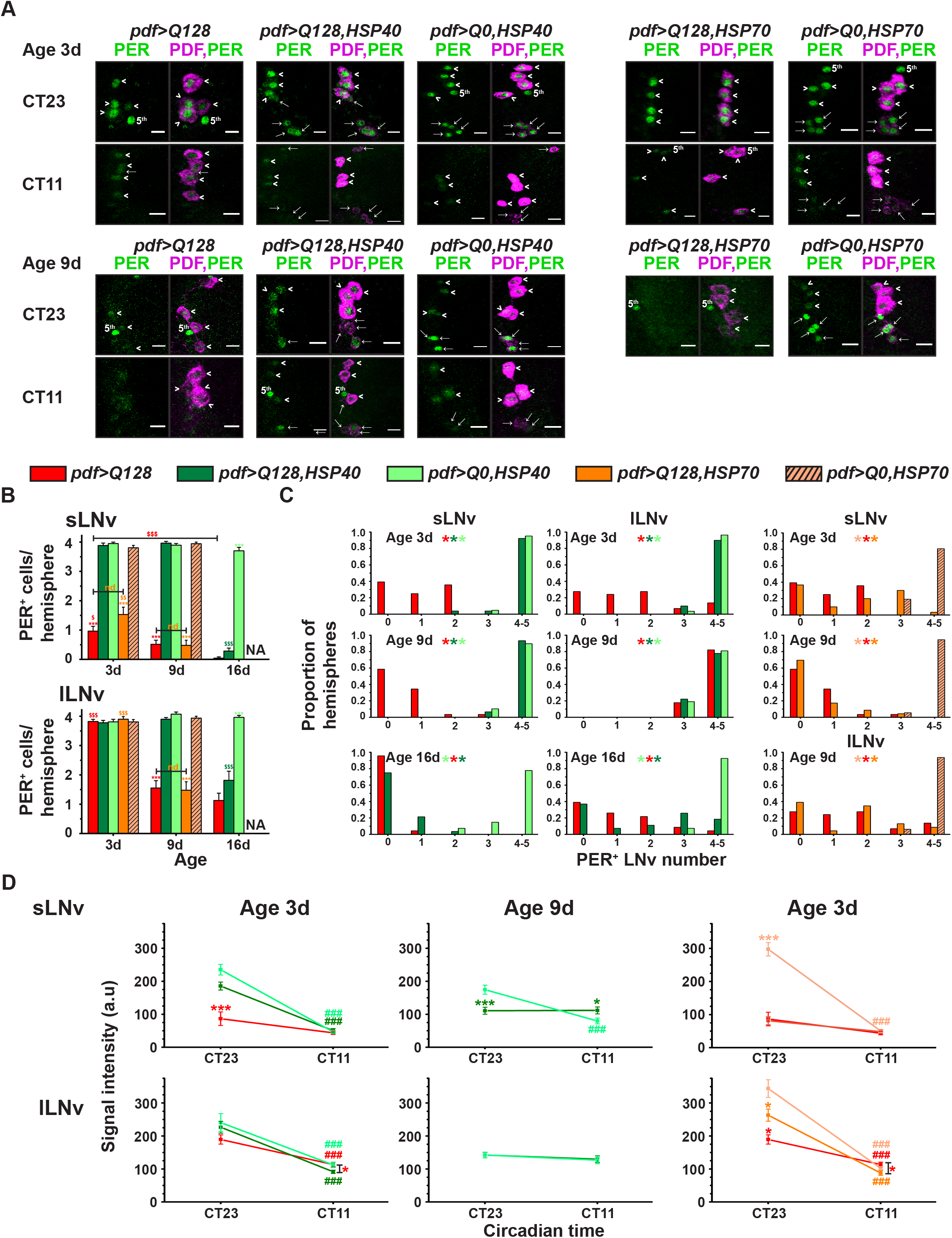
Young *pdf>Q128* flies co-expressing HSP40 show PER oscillations in sLNv. (A) Representative images of adult fly brains stained for PER (green) and PDF (magenta) in LNv at CT23 and CT11. sLNv soma (→ arrows), lLNv soma (> arrowheads) and PDF^-^ PER^+^ 5^th^ sLNv are indicated. Top panel-sets: 3d old flies of five genotypes. Bottom panel-sets: first three panel-sets are of 9d old flies of *pdf>Q128*, *pdf>Q128,HSP40* and *pdf>Q0,HSP40* at CT23 and CT11; fourth- and fifth panel-sets of 9d old *pdf>Q128,HSP70* and *pdf>Q0,HSP70* at CT23. Scale bars are 10 µm. (B) Mean number of PER^+^ sLNv soma (top) and lLNv soma (bottom) at three ages at CT23. Symbols indicate significant differences, * for age-matched, inter-genotype differences and $ for differences between age for each genotype: * (red) of *pdf>Q128* from all other genotypes except *pdf>Q128,HSP70* and * (orange) of *pdf>Q128,HSP70* from other genotypes except *pdf>Q128*. nd is not different. NA is not- applicable. (C) Frequency distribution of the proportion of hemispheres having 0 to 5 PER^+^ LNv soma: of sLNv soma (left) and lLNv soma (middle) in *pdf>Q128,HSP40*, *pdf>Q128* and *pdf>Q0,HSP40* at 3d, 9d and 16d, of sLNv soma at 3d (right-top) and 9d (right-middle) and lLNv soma at 9d (right-bottom) in *pdf>Q128,HSP70*, *pdf>Q128* and *pdf>Q0,HSP70*. Coloured multiple * indicates significantly differing shapes of distribution between genotypes, with the first colour of the reference genotype and the subsequent colours of genotypes differing from the reference at *p*<0.01. (D) Quantification of PER intensity at CT23 and CT11 in sLNv (top) and lLNv (bottom) comparing *pdf>Q128* with *pdf>Q128,HSP40* and *pdf>Q0,HSP40* at 3d (left) and 9d (middle) and comparing *pdf>Q128* with *pdf>Q128,HSP70* and *pdf>Q0,HSP70* at 3d (right). Differences between time-points CT23 and CT11 are represented by # (red) *pdf>Q128*, # (olive-green) *pdf>Q128,HSP40*, # (pale-green) *pdf>Q0,HSP40*, # (orange) *pdf>Q128,HSP70* and # (pale-orange) *pdf>Q0,HSP70*. Coloured * represents age-matched differences of respective-coloured genotype from the indicated one or all others. Statistical significance at single-symbol *p*<0.05, double-symbol *p*<0.01, triple- symbol *p*<0.001. Error bars are s.e.m

Since *pdf>Q128,HSP40* showed control-like PER^+^ sLNv soma numbers at 3d and 9d, PER oscillations in LNv were assessed at these ages. At 3d, *pdf>Q128* did not show PER oscillations in sLNv; *pdf>Q128,HSP40* showed a significant oscillation in PER levels like *pdf>Q0,HSP40* (Fig. 7A, top-left panel-sets, D, top-left). The PER intensity in sLNv of *pdf>Q128* was significantly lower than the other two genotypes at CT23. However, at 9d, despite having control-like sLNv numbers and PER in the sLNv, PER oscillation was absent in sLNv of *pdf>Q128,HSP40* with intensity at CT23 being significantly diminished compared to *pdfQ0HSP40* (Fig. 7A, bottom-left panel-sets, D, top-middle). At 3d, PER oscillation was seen in lLNv of *pdf>Q128*, *pdf>Q128,HSP40* and *pdf>Q0,HSP40* (Fig. 7A, top-left panel-sets, D, bottom-left). At 9d, PER oscillations were absent from lLNv of both *pdf>Q128,HSP40* and control *pdf>Q0,HSP40* (Fig. 7A, bottom-left panel-sets, D, bottom-middle), as is reported for wildtype flies (Shafer et al., 2002; Veleri et al., 2003). PER oscillations in LNv of *pdf>Q128,HSP70* were like *pdf>Q128*, with no PER oscillation in sLNv at 3d, and a significant oscillation in lLNv (Fig. 7A, top-right panel-sets, D, right). HSP70 overexpression did not rescue PER in sLNv, even in young *pdf>Q128* flies. Thus, in young expHTT expressing flies, HSP40 overexpression restores both PER numbers and PER oscillations in the sLNv. This study is the first report thus far of a rescue in circadian molecular oscillations accompanying the restoration of behavioural rhythms observed in these flies, underscoring the effectiveness of HSP40 as a potent circadian modifier in HD.

HSP40 overexpression in LNv of expHTT flies leads to the rescue of sLNv circadian clock output, molecular oscillations, and their associated behavioural rhythms in young flies. The rescue in behaviour and the clock output PDF is long-lasting. There is also a considerable reduction in the expHTT inclusion load. This sustained rescue at multiple levels posits HSP40 as a potential therapeutic candidate in improving circadian health under neurodegenerative conditions.

## Discussion

### HSPs as modifiers of HD-induced circadian dysfunction

The role of HSPs, known modifiers of neurodegeneration, in HD-associated circadian disturbances is relatively unexplored. In this study, we show that the overexpression of the co-chaperone HSP40 in circadian pacemaker neurons of *Drosophila* alleviates expHTT- induced circadian behavioural arrhythmicity and suppresses circadian neurotoxicity. Overexpression of the central chaperone HSP70 also mitigates expHTT-induced circadian behavioural rhythm disruptions, but not cellular phenotypes. The rescue upon HSP40 overexpression was more robust, pronounced and sustained. In young flies, HSP40 overexpression seems to restore the functionality of LNv, particularly the central pacemaker sLNv. Evidence for sLNv functionality is the restoration of circadian proteins, the core-clock protein PER oscillations and output neuropeptide PDF in the sLNv soma and lowered expHTT inclusion load, leading to an overall improvement in the LNv-circuit associated behavioural rhythms. These flies continued to be behaviourally rhythmic up to 16d but with lowered robustness, with the presence of PER in LNv and control-like PDF^+^ sLNv soma numbers and diminished inclusion load, but without PER oscillations. The persistence of activity rhythms in the absence of PER oscillations in sLNv indicates that these oscillations might be dispensable for rhythm sustenance. Two recent studies that support this reasoning show that PER in LNv does not seem necessary for the persistence of free-running activity rhythms but is vital for rhythm strength (Delventhal et al., 2019; Schlichting et al., 2019). In relatively older flies, despite the presence of nearly 4 PDF^+^ sLNv soma (16d) and reduction in the inclusion load (aggregate numbers of 16d old *pdf>Q128,HSP40* flies were comparable to 9d; quantification not shown), the flies were arrhythmic during AW3 (16d-23d). Thus, the HSP40 neuroprotection does not seem sufficient as the flies age, contributing to a deterioration of LNv health. The inadequacy of HSP40 expression to extend protection to LNv for extended durations suggests two conclusions. Firstly, the restoration of PDF^+^ sLNv does not guarantee sustained free-running rhythms in the absence of PER. Our previous results show that about 20% of 7d old arrhythmic *pdf>Q128* flies had at least 1-2 PDF^+^ sLNv. The PDF levels in sLNv dorsal projections were oscillating and functional in synchronising downstream circadian neurons (Prakash et al., 2017). The previous results, together with our observations of behavioural arrhythmicity in older flies despite PDF rescue in sLNv, indicates that sLNv PDF, in the absence of PER, is insufficient for rhythmic activity. Secondly, over time, neuroprotective benefits offered by HSP40 can be overwhelmed upon HD progression, probably by the age-related burden on LNv proteostasis, rendering the cells vulnerable to expHTT toxicity. Therefore, sustained rhythm rescue might require supplementing HSPs with further enhancement of proteostasis via proteasomal or autophagic upregulation.

HSP70 overexpression, in contrast, showed a rescue in only early-age rhythms and activity consolidation, albeit of lowered robustness, a decrease in early-age inclusion number, without the rescue of PDF^+^ sLNv numbers or PER oscillations in sLNv or alterations to inclusions being the prevalent form of expHTT in LNv. The rhythmic flies of *pdf>Q128,HSP70* have poor rhythm robustness and can be attributed to the absence of PER rescue in the LNv. However, the persistence of behavioural rhythms on HSP70 overexpression in the absence of PDF rescue in the sLNv soma is intriguing. It suggests that the presence of PDF in sLNv is not necessary for behavioural rhythm restoration. Other studies in *pdf>Q128* have reported rhythm rescue with only marginal PDF restoration in sLNv soma upon ATX2 or HOP down- regulation (Xu et al., 2019b; Xu et al., 2019a). Together with ours, these reports suggest that other mechanisms might drive circadian behavioural rhythms without canonical circadian cellular proteins. Possible intersections of HSP onto LNv function and output in orchestrating rhythmicity are circadian oscillations in arborisations of sLNv termini and secondary molecular loop components, non-PER driven clocks, LNv membrane properties, neuronal firing, synaptic strength and network-level communication (Edgar et al., 2012; Beckwith and Ceriani, 2015; Yao et al., 2016; Rey et al., 2018; Bulthuis et al., 2019).

### Impact of HSP overexpression on visible inclusions of expHTT

HSPs dilute the presence of aggregate-prone proteins by interfering with the aggregation pathway like delaying nucleation, fibril elongation or redirecting the pathway towards less-toxic versions, sequestration into cellular compartments or organelles and targeting for degradation (Kampinga and Bergink, 2016; Mannini and Chiti, 2017; Hipp et al., 2019). The effect of HSPs on aggregation is variable. It depends on a host of factors like the definition of aggregates, their nature and conformation, the cellular context, the stage of aggregation, age and disease stage, quantification method, and model system. Indeed, with up-regulation of HSPs (HSP40 and HSP70) in HD models, there is evidence for differential effects: many showing a decline in aggregation (Jana, 2000; Zhou et al., 2001; Hay, 2004; Guzhova et al., 2011; Labbadia et al., 2012; Maheshwari et al., 2014; Scior et al., 2018), some showing no effect (Wyttenbach et al., 2000; Karpuj et al., 2002; McLear et al., 2008) and one study showing increased aggregation (Wyttenbach et al., 2000).

In the present study, the visible puncta-like clumped appearance of HTT-Q128 as detected under a fluorescence light microscope using immunocytochemistry is defined as an inclusion. This definition excludes detecting many species below the resolution limit and does not distinguish based on solubility and other biochemical features. Hence, the inferences are limited to the size range and gross features of particles detected via an epifluorescence scope. However, this does not take away the validity of the effect of HSP40 overexpression on expHTT inclusions and its impact on LNv function at the cellular and behavioural stages. In *pdf>Q128* flies, HSP40 overexpression in the LNv leads to a reduction in the number of visible expHTT inclusions and a decline in the dominance of inclusion form of expHTT with the appearance and dominance of expHTT spots. Accompanying them are improvements in LNv pacemaker function as evidenced by the re-establishment of circadian molecular and behavioural rhythms. Thus, a decreased inclusion load and the dominance of expHTT spots could lead to enhanced functionality of the LNv.

A pictorial representation of the locomotor behaviour and cellular phenotypes in LNv comparing *pdf>Q128* with *pdf>Q128,HSP40* is shown (Fig. 8). A clear pattern for expHTT forms in LNv with age emerges. For the toxic *pdf>Q128* and relatively less-toxic *pdf>Q128,HSP70*, Diff+Inc expHTT in lLNv at an early age gives way to inclusions-only at later ages. In the neuroprotective *pdf>Q128,HSP40*, across age, Spots are present in sLNv, dominating over Inc at 3d, while Inc dominates at later ages. In the lLNv of *pdf>Q128,HSP40*, Diff expHTT dominates at 3d, giving way to a combination of diffuse+spots and inclusions at 9d, and then to non-diffuse expHTT (Spot and Spot+Inc) at 16d. In *pdf>Q128,HSP40*, the continued presence and domination of expHTT Spot in LNv is associated with intact PDF^+^ sLNv and behavioural rhythmicity of most of the *pdf>Q128,HSP40* flies up to 16d. Together, these results indicate an association between the expHTT form predominating in LNv and LNv health, that of diffuse and spot forms with healthy LNv and rhythmic activity, while inclusions with poor LNv function and arrhythmicity.

**Fig. 8.**
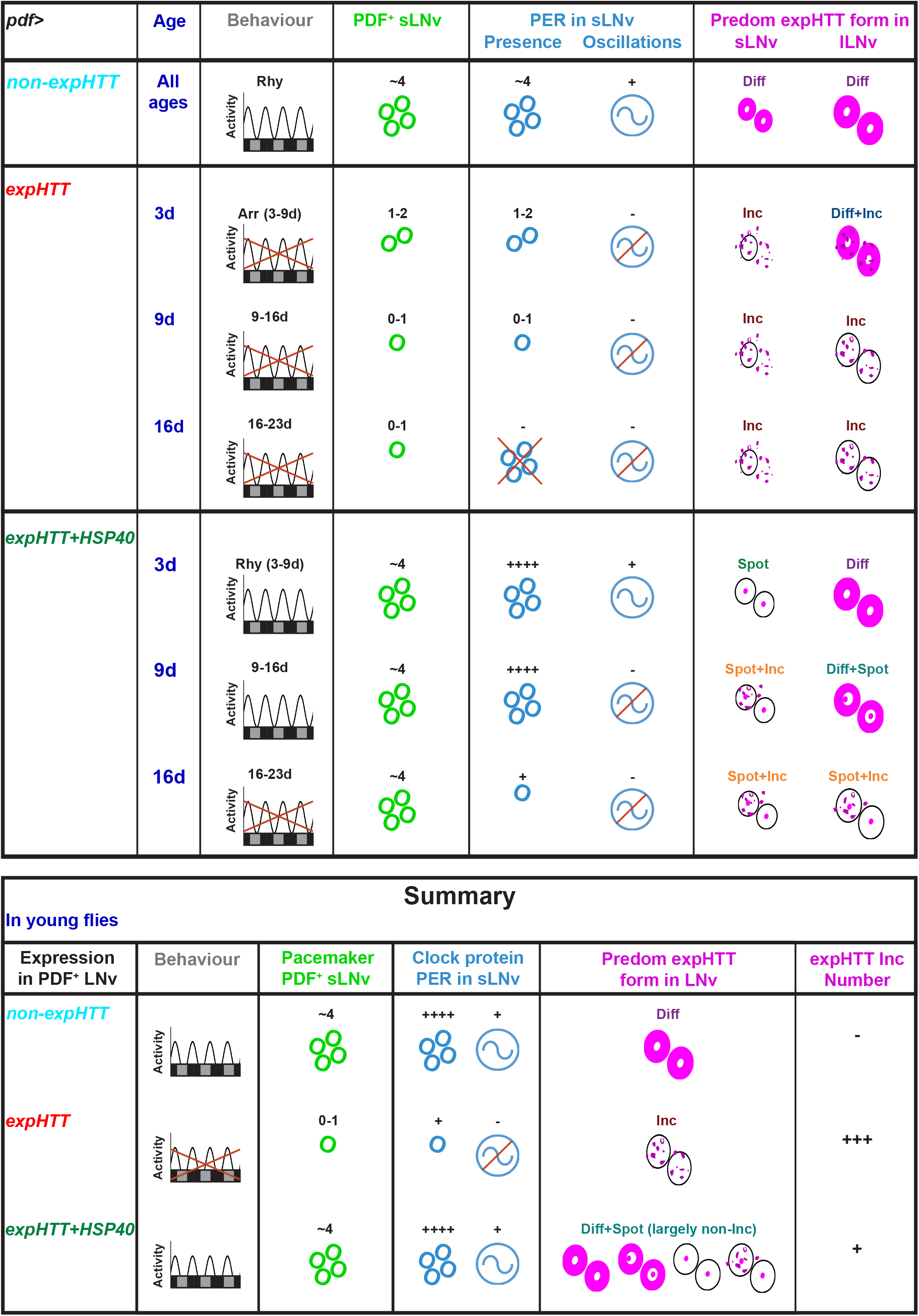
HSP40 is neuroprotective and delays circadian dysfunction in HD: A graphical summary. Top table: A pictorial representation of the effect of expressing expHTT alone and with HSP40 in the LNv of *Drosophila* on circadian neurodegenerative phenotypes across age. The control phenotype on expressing non-expanded HTT (Q0) in LNv is shown at the top. The effects on the circadian behavioural activity/rest rhythms, PDF^+^ and PER^+^ sLNv soma numbers, PER oscillations in sLNv, and predominant form of expHTT in sLNv and lLNv are shown across age. The behavioural rhythms are represented for three 7d age windows, whereas the cellular phenotypes are for specific ages. Arr stands for arrhythmic and Rhy for rhythmic. Bottom table: A summary of the key findings. Co-expressing HSP40 with expHTT in the LNv reverses the expHTT-induced circadian phenotypes of behavioural arrhythmicity, PDF loss from sLNv soma and loss of PER oscillations and PER in sLNv of young flies. Also, the expHTT inclusions, a characteristic neurodegenerative phenotype, and the predominant expHTT form observed in the LNv of *pdf>Q128* flies are replaced by mainly non-inclusion forms: diffuse, spots and a combination. The prevalence of non-Inc expHTT forms is also reflected as a decrease in the number of expHTT inclusions. In summary, HSP40 is an effective suppressor of HD-induced circadian disruptions

HSP70 overexpression in *pdf>Q128* flies rescues early-age rhythms, reduces expHTT inclusion numbers, but with the dominance of inclusion as the main expHTT form in LNv, suggesting that HSP70-mediated improvements to LNv health are via inclusion-independent mechanisms. HSP70 serves aggregation-independent neuroprotective roles like inhibiting apoptosis (Beere, 2004; Kennedy et al., 2014), combating inflammation (Borges et al., 2012; Dukay et al., 2019), reducing ROS and oxidative stress (Kalmar and Greensmith, 2009; Wyttenbach and Arrigo, 2009) and supporting synaptic function (Deane and Brown, 2016; Gorenberg and Chandra, 2017). Such aggregation-independent neuroprotection by HSP40 and HSP70 in HD is reported (hou et al., 2001; Wyttenbach, 2002; Borrell-Pages et al., 2006; Wacker et al., 2009). The versatility of HSP70 in enhancing neuronal health and function could contribute to circadian rhythm improvements in the absence of an effect on inclusion load and PDF restoration in sLNv.

### Spot form of expHTT

Upon overexpression of HSP40 in *pdf>Q128* flies, a new form of expHTT with a spot-like appearance located close to the nucleus and seemingly overlapping with the nuclear PER was observed. We discuss the possible significance of the spot form of HTT. In eukaryotes, aggregate-prone proteins are often sequestered into specialised cellular compartments that are thought to be neuroprotective, are typically membrane-less, sometimes referred to as sequestrosomes or into membrane-bound organelles (Sontag et al., 2014; Tan and Wong, 2017; Johnston and Samant, 2021). A few examples of such spatially-sequestered quality control sites are the cytoplasmic Q-bodies or stress foci, cytoplasmic p62 bodies or ALIS (aggresome-like induced structures), peri-nuclear JUNQ (Juxta Nuclear Quality Control Compartment), intra-nuclear INQ (intranuclear quality control compartment), peri-nuclear aggresomes and peri-vacuolar IPOD (insoluble protein deposit) (Tan and Wong, 2017; Johnston and Samant, 2021). HSPs are known to participate in such compartmentalisations (Nollen et al., 2001; Specht et al., 2011; Escusa-Toret et al., 2013; Miller et al., 2015). The aggresomes are mostly juxta-nuclear, membrane-free inclusions carrying ubiquitinated misfolded proteins formed at the microtubule organising centre (MTOC) (Johnston et al., 1998; Kopito, 2000; Johnston and Samant, 2021). There is evidence for colocalization of HSP40, and HSP70 colocalise with aggresomes (Garcia-Mata et al., 1999; Junn et al., 2002; Gamerdinger et al., 2011; Zhang and Qian, 2011).

Another observation here is that the average size of an expHTT Spot in the small LNv is significantly larger than those in the large LNv. This sizeable expHTT Spot in the sLNv could reflect a greater expHTT burden and toxicity in the vulnerable sLNv or a longer duration of HTT and HSP40 expression owing to their earlier appearance than lLNv during development (Helfrich-Forster, 1997). The presence of 3-4 PDF+ sLNv in nearly every hemisphere of *pdf>Q128,HSP40* across age, parallels with the presence of at least one sLNv per hemisphere having expHTT Spot (proportion of sLNv per hemisphere with at least one expHTT Spot: 3d-100%, 9d ∼ 97% and 16d ∼79%). Such co-occurrences indicate that the appearance of spotted expHTT might be protective. Thus, HSP40 might modify the nature of expHTT inclusions, and the spotted expHTT could represent a less reactive and relatively benign form of expHTT, contributing to an enhancement in LNv health and function.

### Effect of HSP40 and HSP70 co-expression

Many studies in PolyQ models have described a synergistic effect of co-expression of HSP40 and HSP70 toxicity: expression of both offered better protection than either alone (Chan et al., 2000; Jana, 2000; Kobayashi et al., 2000; Muchowski et al., 2000 Sittler et al., 2001 Bailey et al., 2002; Bonini, 2002; Rujano et al., 2007). The current study shows a synergistic effect of improving the HD-induced daily circadian activity consolidation in young flies, but not on rhythmicity and rhythm strength. The synergistic effect seems subtle, evident with a daily readout like ‘*r*’, but not with a 7d-overt-readout of rhythmicity. Over time, there was a decline in rhythm robustness of *pdf>Q128* flies expressing both HSPs than those expressing HSP40 alone, suggesting that co-expression of multiple HSPs could become detrimental over time. In another study, such co-expression eliminated the survival benefit of HSP70-only expression, enhancing cell death (Ormsby et al., 2013). Some of the drawbacks of HSP co-expression and HSP overexpression, like the pro-carcinogenic effect and generation of seeding-competent nuclei, call for caution when targeting central HSPs for therapy (Jaattela, 1995; Nylandsted et al., 2002; Tittelmeier et al., 2020b).

### HSP40 vs HSP70: HSP40, a superior suppressor of HD neurotoxicity?

In our study, HSP40 emerged as a superior suppressor of most of the expHTT-induced phenotypes examined. There is substantial support for HSP40 being a more effective HD neurotoxic modifier. Among the HSP40, HSP70 and HSP110 chaperone families, the DNAJB class of HSP40 emerged as the most potent protector against PolyQ toxicity (Hageman et al., 2010), and in R6/2 mice, HSP70 suppressed HD only moderately (Hansson et al., 2003; Hay, 2004; Popiel et al., 2012), while HSP40 members had better success (Labbadia et al., 2012; Kakkar et al., 2016). HSP40 also prevented the secretion of expanded PolyQ proteins from cultured cells and can likely prevent cell-to-cell transmission (Popiel et al., 2012), an emerging concern in NDs. Findings from the present study and other studies (Chai et al., 1999; Zhou et al., 2001; Rujano et al., 2007; Ormsby et al., 2013) show that HSP40 reduces aggregation more often than HSP70. HSP40 is rate-limiting in the suppression and reversal of expHTT aggregation by disaggregases (Rujano et al., 2007; Scior et al., 2018), and some members can act without requiring HSP70 (Hageman et al., 2010; Kuo et al., 2013; Månsson et al., 2013; Kakkar et al., 2016). The significant role of the DNAJB protein family in synaptic health and neuronal proteostasis, their diversity in function, distribution and substrate specificity, underscores their usefulness in directed therapy while minimising side-effects (Chuang et al., 2002; Westhoff et al., 2005; Gibbs et al., 2009; Kampinga and Craig, 2010; Gao et al., 2015; Nillegoda et al., 2018; Kampinga et al., 2019; Tittelmeier et al., 2020a).

### A need for screening circadian-specific neurotoxic modulators

In an *in vivo* system, we have shown a neuroprotective role of chaperone HSP40 in rescuing HD-induced circadian deficits and neurotoxicity at multiple levels and across the temporal scale. Such a multi-level associated functional rescue offers an edge over conventional non-associated cellular and behavioural rescues like making better cause-effect inferences due to reduced off-target effects, rigorous testing of the robustness and versatility of the modifying treatment, and serving as proof-of-principle evaluations. Also, though most of the candidates screened are known modulators of neurotoxicity, only two groups of proteins emerged as potent suppressors of circadian disruption, uncovering a gap in establishing circadian-specific neuroprotective agents.

### HSPs, circadian health and neurodegenerative diseases

There is ample evidence for clock-control regulation of proteostasis components, including chaperones (Desvergne and Friguet, 2017; Ryzhikov et al., 2019; Wang et al., 2020), with both HSP40 and HSP70 isoforms, showing rhythmic gene expression across taxa (Li et al., 2017). The converse, i.e. proteostasis affecting molecular clock via post-translational modifications and autophagy, is also prevalent (Mehra et al., 2009; Stojkovic et al., 2014; Toledo et al., 2018; Juste et al., 2021). However, very few studies have assessed the role of HSPs in circadian maintenance and its deterioration in NDs, especially in animals. In *Drosophila*, the Hsp70/Hsp90-organizing protein (HOP) improved rhythmicity in an HD model (Xu et al., 2019b), HSP70 expression overcame arrhythmicity on Gal4-overexpression in the LNv (Rezaval et al., 2007), and HSPs indirectly implicated in circadian behaviour (Benbahouche Nel et al., 2014; Means et al., 2015). In mouse fibroblasts, HSP90 is required for circadian rhythmicity, while HSF1 and ER HSP70 strengthened rhythms post-stress (Tamaru et al., 2011; Schneider et al., 2014; Pickard et al., 2019). The present findings of a relatively novel role for the HSPs in protecting against ND-induced circadian dysfunction and the above studies encourage further research on HSPs in circadian function. Given that circadian disturbances are early and pre-manifest in HD, treatments targeting HSPs could impact the early stages of HD and have a meaningful therapeutic impact. With an ageing population worldwide, there is an increasing prevalence of NDs (Gitler et al., 2017; Lassonde, 2017; Bejot and Yaffe, 2019). Given the pivotal roles of proteostasis and circadian health in NDs, studying the involvement of molecular chaperones in circadian maintenance will significantly improve our understanding of ND progression and treatment.

## Acknowledgements

We sincerely thank Troy Littleton, Massachusetts Institute of Technology, USA, Michael Nitabach, Yale University, USA, Jeffrey C Hall, Brandeis University, USA and Jae Park, Vanderbilt University, USA, for sharing reagents. Special thanks to Florence Maschat, Université de Montpellier, France for the *UAS-dHTT81aa* and *UAS-dHTT620aa* lines (Mugat et al., 2008) and Norbert Perrimon, Harvard Medical School, USA for the *UAS-dHTT* line (Zhang et al., 2009). We thank Sunil Kumar from the JNCASR Confocal Imaging Facility for his assistance. We thank Ankit Sharma for his comments on the manuscript and help with experiments and Rutvij Kulkarni and Abhilash Lakshman for statistical analyses. We thank Sushma Rao for help with dissections and Rajanna and Muniraju for their assistance.

## Footnotes

### Author contributions

Conceptualization: P.P., V.S.; Methodology: P.P., V.S.; Validation: P.P., Formal Analysis: P.P., A.K.P.; Investigation: P.P., A.K.P.; Resources: V.S.; Writing - original draft: P.P.; Writing- review and editing: P.P., A.K.P., V.S.; Visualisation: P.P., V.S.; Supervision: V.S.; Project administration: V.S.; Funding acquisition: V.S.

### Data availability

Upon acceptance raw data will be placed in the DSpace repository at http://lib.jncasr.ac.in:8080/xmlui/handle/

